# Temporal Coding rather than Circuit Wiring allows Hippocampal CA3 Neurons to Dynamically Distinguish Different Cortical Inputs

**DOI:** 10.64898/2026.02.20.707083

**Authors:** Keelin O’Neil, Vincent Robert, Luke A. Arend, Buyong Kim, André A. Fenton, Jayeeta Basu

## Abstract

Memory relies on associating and indexing multimodal information. How does this occur within single neurons? The hippocampus integrates multisensory information from medial (MEC) and lateral (LEC) entorhinal cortices to form environmental representations^1-6^, yet their synaptic dynamics, circuit organization, and integrative functions remain elusive. Contrary to canonical models emphasizing anatomical and functional segregation, our dual-color optogenetic^7^ circuit mapping revealed that both MEC and LEC inputs converge on virtually every CA3 pyramidal neuron and exhibit similar input-output functions and modulation via GABAergic microcircuitry. Divergence occurs in their frequency-dependent short-term plasticity. With increasing frequencies, facilitating LEC-evoked responses undergo synaptic depression, whereas MEC-evoked responses continue facilitating. *In vivo,* MEC-originating dentate spikes fire CA3 more than LEC, likely due to their distinct temporal relationship with oscillatory activity. Our findings support that hippocampal area CA3 processes region-specific information from MEC and LEC via temporal coding, rather than hard-wired circuit organization, at both single-neuron and network levels.

## Introduction

A fundamental question in neuroscience is how the brain processes multimodal information in a versatile and efficient manner while maintaining content specificity. Principal neurons across the brain integrate multiple inputs from various brain regions, conveying information about distinct aspects of the external environment, experiences, and internal states^4,8,9^. Both integration and segregation of information could be implemented anatomically, via hard-wired circuits, or temporally, via activity-dependent dynamic states of individual neurons and networks^10,11^. To parse how such information multiplexing is achieved, we turned to the cortico-hippocampal circuit model to explore how single neurons distinguish and associate different types of information shaping relevant network dynamics.

Pyramidal neurons (PNs) in the hippocampus receive spatial information from the medial entorhinal cortex (MEC) and sensory-contextual information from the lateral entorhinal cortex (LEC). Canonically, MEC and LEC have been studied and treated as independent coding structures with anatomical and functional segregation^8,12-15^. MEC harbors grid cells^14,16^, border cells^17,18^, and head-direction cells^19,20^ that project space-related activity to HC^16,21,22^. Despite the hypothesized relationship between spatially tuned MEC inputs and hippocampal place systems, inactivation of MEC has a modest impact on hippocampal spatial coding and related behaviors^23-29^. This suggests additional inputs are involved in encoding and maintaining these representations. On the other hand, LEC encodes contextual features such as objects and their displacement^30-32^, odors^33-35^, and salient rewarding^36,37^ or aversive stimuli^38-41^ to support complex learning rules and temporal sequences^38,42-44^. Nevertheless, recent studies have shown pronounced effects of LEC inactivation on remapping^45^ and learning-driven stabilization of CA3 place cells^46^. These findings, along with other studies^47-49^, suggest that the association of MEC and LEC inputs within the hippocampus is likely a crucial step in forming multimodal episodic memories for generating adaptive learned behaviors^3,25,45,49-51^. However, circuit interactions between MEC and LEC have rarely been examined within the same animal, at either the single-cell or network level.

CA3 pyramidal neurons (PNs) embedded within an auto-associative recurrent network provide an ideal model system to study this problem^52-58^. Unlike in CA1, where the MEC and LEC inputs are anatomically segregated along the proximal-distal axis^59,60^ or in the Dentate Gyrus (DG), where these inputs are clearly stratified within dendritic compartments^61-63^, in CA3, inputs from layer II of MEC and LEC are not as compartmentally distinct. In addition, CA3 PNs receive indirect MEC and LEC information routed through the DG and CA2 circuits^15,46,57,60,64^. By virtue of this skip connection^65^, contextual and spatial inputs can be directly bound within a single CA3 cell through dendritic integration. This direct input serves as an instructive teaching signal to improve CA3 processing of incoming indirect information from MEC and LEC^66-68^, hierarchically integrated at the level of DG^11,61-63^ or cortex itself ^22,59^. While discrimination of cortical information in the hippocampal system has been observed *in vivo* at the network level through temporally distinct oscillatory synchrony^10^ with MEC (fast gamma) ^49,51,69^ and LEC (slow gamma) ^51,70^, we know little about how single neurons implement this through synaptic and circuit interactions.

To understand whether CA3 processing of multisensory information uses temporal coding or circuit organization (single-neuron or network-level), we explored the biophysical and microcircuit organization of glutamatergic inputs to CA3 from LEC (LEC_glu_) and MEC (MEC_glu_) as well as the *in vivo* network discharge dynamics driven by these two inputs. We used dual-color optogenetics and patch-clamp electrophysiology-based functional circuit mapping *ex vivo,* as well as large-scale extracellular electrophysiology *in vivo*, to physiologically dissect the MEC and LEC glutamatergic projections to CA3 circuit and resultant network activity. Our findings suggest the wiring and output of these circuits are largely similar at the level of single neurons, with two exceptions: their activity-dependent temporal dynamics and their monosynaptic excitatory drive onto CA3 PNs. Comparing *in vivo* CA3 responses to spontaneous dentate spikes originating from either MEC (DS_MEC_) or LEC (DS_LEC_) reveals that CA3 network activity is largely driven by MEC, irrespective of the order and timing between DS_LEC_ and DS_MEC_. Our findings support a growing literature showing that the relative timing of these two distinct inputs is critical for conveying information and that temporal shifts can amplify small differences in neuronal activation to yield network-wide effects.

## Results

### Individual CA3 pyramidal neurons receive monosynaptic connections from both MEC & LEC inputs

MEC and LEC send glutamatergic projections to the distal dendrites of hippocampal area CA3^5,57,71^, but whether they form synapses on the same CA3 pyramidal neuron (PN) remains unknown. To map the MEC_glu_ and LEC_glu_ projections to CA3 in the same samples, we injected two adeno-associated viral (AAV) constructs encoding Cre-dependent spectrally separated opsins (ChrimsonR and Chronos)^7,46^ in MEC and LEC of a CaMKII-Cre mouse line which selectively expresses Cre in glutamatergic neurons^46^ (Fig. 1a). Our stereotaxic injections successfully restricted the viral expression to the injected regions (Fig. 1b). We observed MEC_glu_ and LEC_glu_ axons in the *stratum lacunosum moleculare* (SLM) layer of CA3, with LEC_glu_ axons more distal from the pyramidal layer (*stratum pyramidale*, SP) than MEC_glu_ axons (Fig. 1b-d, Extended Data Fig. 1a-c). In addition, we observed the same innervation patterns in separately (single) labelled animals (Fig. 2i,j). LEC_glu_ projections occupy more area in the SLM region of CA3 than MEC_glu_ in our preparation (Extended Data Fig 1. b,c), consistent with previous reports^55-57^.

**Figure 1:**
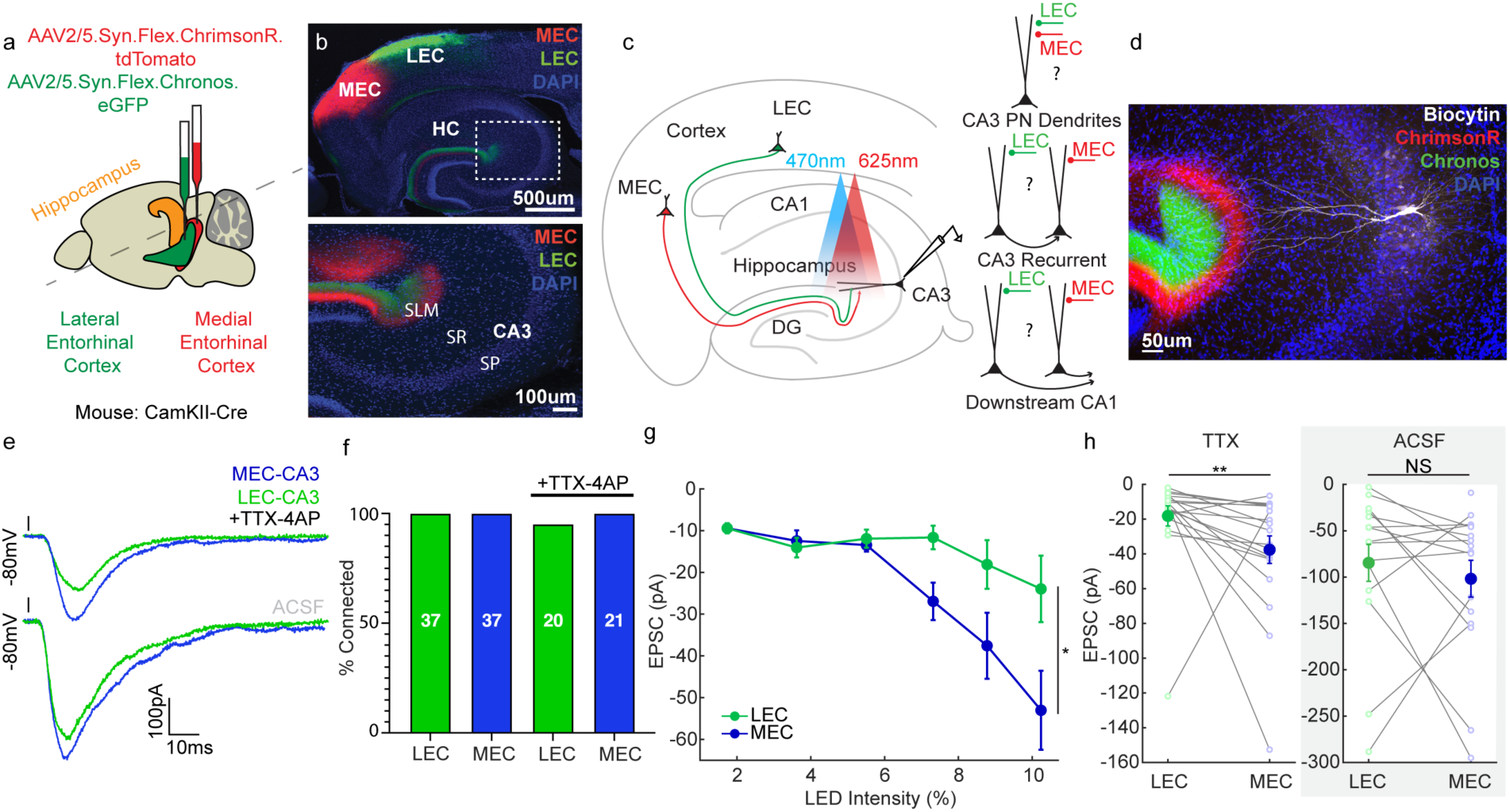
Individual CA3 PNs receive monosynaptic inputs from both MEC_glu_ and LEC_glu_. **a,** Schematic of viral injection strategy in MEC and LEC. Sagittal view of mouse brain showing MEC (red), LEC (green), and Hippocampus (orange). The dotted line represents the plane of the section. AAV2/5.Syn.Flex.ChrimsonR.tdTomato and AAV2/5.Syn.Flex.Chronos.eGFP were injected into MEC and LEC of CaMKII-Cre mice to drive excitatory opsin expression selectively in glutamatergic neurons. We alternated the opsins between the two regions to avoid confounds due to opsin-specific effects. This enabled input-specific dual-optogenetic circuit mapping of MEC and LEC inputs to hippocampal area CA3. **b,** (top) Representative confocal images of injection sites in a 100 µm horizontal section through the dorsal hippocampus and entorhinal cortex showing restricted expression of fluorophores in MEC and LEC. This brain slice was stained for DAPI, a nuclear marker. (bottom) Expanded view of the area enclosed in the dotted box in the top image shows axonal expression of fluorophores in *stratum lacunosum moleculare* (SLM) of CA3. MEC and LEC projections show typical patterning, MEC axons more proximal to stratum pyramidale (SP) than LEC axons. **c,** (left) Input-specific, dual-optogenetic functional circuit mapping in hippocampal area CA3 in acute mouse brain slice experimental design. Optogenetic stimulation of MEC_glu_ and LEC_glu_ axons via 470 nm and 625 nm light while performing whole-cell voltage-clamp recordings of CA3 pyramidal neurons (PN). (right) Schematic of connectivity hypothesis. **d,** Confocal image of CA3 region of 400 µm transverse hippocampal, mouse brain slice showing recorded CA3 a/b PN filled with Biocytin (white) and MEC (red) and LEC axons (green). The slice is stained with DAPI and streptavidin-AF647 (Biocytin counterstain). **e,** Sample traces of MEC_glu_-(blue) and LEC_glu_-driven (green) responses in CA3 PN voltage-clamped at −80 mV in 1 µM TTX & 10 µM 4-AP (top) and ACSF (bottom). **f,** Fraction of CA3 PN that receives MEC_glu_ (blue) and LEC_glu_ (green) inputs in ACSF and TTX & 4-AP. **g,** Input-output function curves of MEC_glu_ (blue) and LEC_glu_ (green) light-evoked CA3 PN EPSCs in TTX & 4-AP across low (<10%, 0.31 mW/mm^2^) light intensities (n = 19 neurons, N = 10 mice, *p = 0.0429 Mixed-Effects Model w/ Geisser-Greenhouse Correction). **h,** Peak amplitudes of LEC_glu_ (green) and MEC_glu_ (blue) light-evoked CA3 PN EPSC responses at 8.8% light intensity in TTX & 4-AP (left, n = 19 neurons, N = 10 mice) and ACSF (right, n = 15 neurons, N = 8 mice) (TTX & 4-AP **p = 0.0039 Wilcoxon matched-pairs two-tailed,, ACSF NS p = 0.3591 Wilcoxon matched-pairs two-tailed). Each data point represents the mean peak EPSC amplitude of an individual cell. All error bars represent SEM.

**Figure 2:**
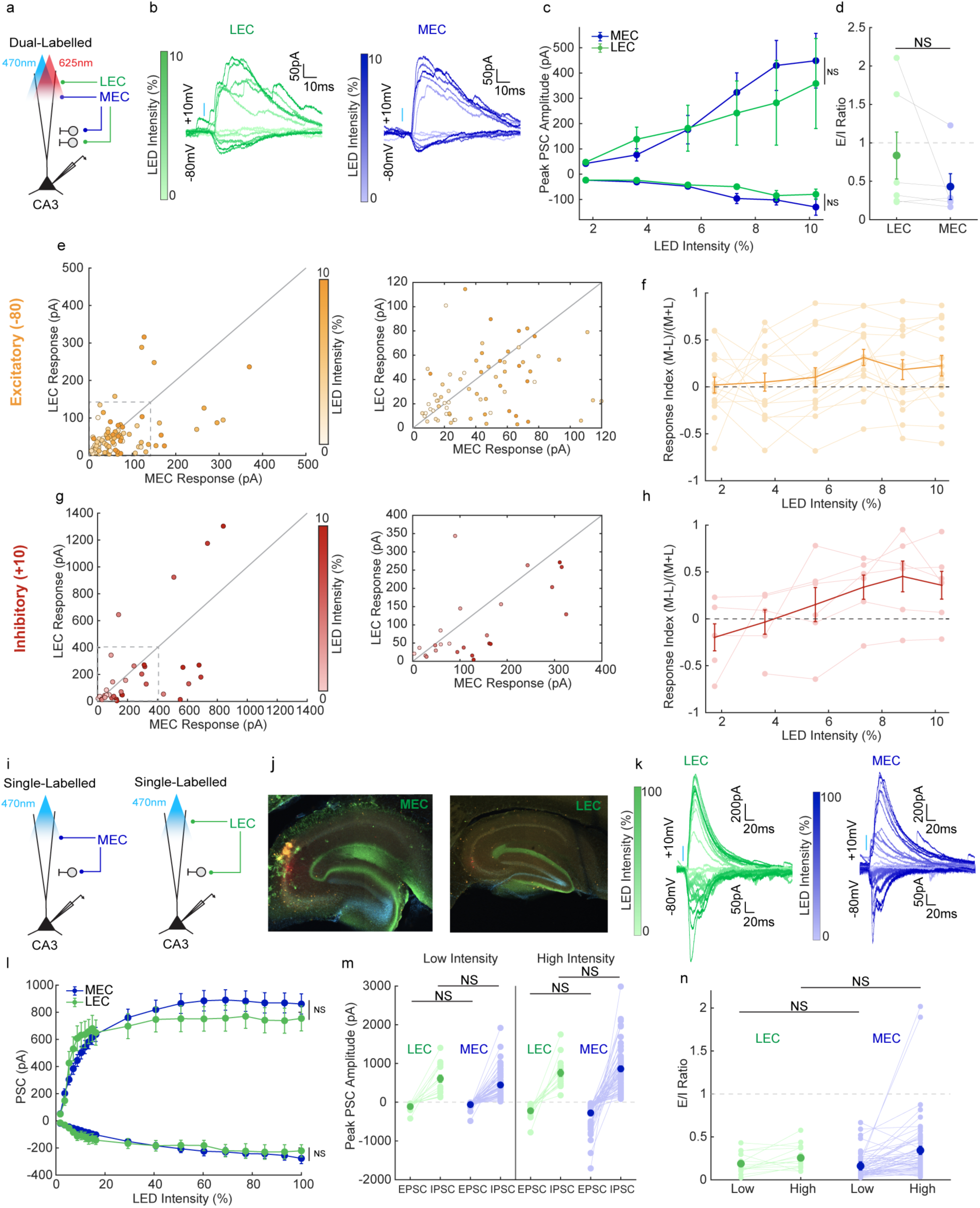
MEC_glu_ and LEC_glu_ driven excitation and inhibition in CA3 PN is similar in both putative monosynaptic and polysynaptic regimes. **a,** Dual-labelled experimental strategy described in Figure 1a-d. We recorded responses of CA3 PN voltage-clamped at +10 mV and −80 mV to LEC_glu_ (green) and MEC_glu_ (blue) stimulation done using 470 nm and 625 nm light over the SLM region of CA3. **b,** Sample traces of LEC_glu_-(green) and MEC_glu_-driven (blue) CA3 PN EPSCs (−80 mV) and IPSCs (+10 mV) at a range, 1-10% (0.05-0.31 mW/mm^2^), of LED intensities. Responses to MEC_glu_ and LEC_glu_ are recorded from the same cell. Saturation of trace indicates LED intensity. **c,** Input-output function curves of MEC_glu_ (blue) and LEC_glu_ (green) light-evoked CA3 PN excitatory (EPSCs) and inhibitory post-synaptic potentials IPSCs across low light intensities (IPSC: n = 6 cells, N = 4 mice, NS p = 0.7960 Mixed-effects model w/ Geisser-Greenhouse Correction, EPSC: n = 16 cells, N = 8 mice, NS p = 0.1332 Mixed-effects model w/ Geisser-Greenhouse Correction). **d,** Excitation/inhibition (E/I) ratios of MEC_glu_-(blue) and LEC_glu_-driven (green) CA3 PN EPSCs/IPSCs. Each data point represents the E/I ratio of an individual neuron and only neurons from which both EPSCs and IPSCs were recorded are included in the E/I ratio analysis. (n = 6 cells, N = 4 mice, NS p = 0.0625 Wilcoxon matched-pairs two-tailed). **e, g,** Averaged individual peak amplitude CA3 PN EPSC (**e,** n = 16 cells, N = 8 mice) and IPSC (**g,** n = 6 cells, N = 4 mice) responses to MEC_glu_ (x-axis) and LEC_glu_ (y-axis) stimulation across range of low LED intensities. Saturation of trace indicates LED intensity. Expanded view of demarcated area shows spread of data points with peak amplitudes from 0pA to 120pA. **f,h,** Response index for MEC_glu_- and LEC_glu_-driven EPSCs (**f,** n = 16 cells, N = 8 mice) and IPSCs (**h,** n = 6 cells, N = 4 mice) in CA3 PN. Each connected dot is an average for an individual cell across low light intensities. **i,** Single-labelled experimental schematic. Mice were injected in either MEC or LEC with AAV2/5.Syn.Flex.Chronos.eGFP viral construct in a CaMKII-Cre mouse or AAV2/5.CaMKII.Chronos.eGFP in a wild-type mouse to drive excitatory opsin expression selectively in glutamatergic neurons. This enabled input specific functional circuit mapping in separate animals to explore a further range of stimulation intensities (1-100% 0.05-3.02 mW/mm^2^). **j,** Confocal images of transverse hippocampal recording slices with MEC_glu_- (left) or LEC_glu_- (right) axons (green) labelled. **k,** Sample traces of LEC_glu_- (green) and MEC_glu_-driven (blue) CA3 PN EPSCs (−80 mV) and IPSCs (+10 mV) at a range, 1-100%, of LED intensities. Responses to MEC_glu_ and LEC_glu_ are from different cells/mice. Saturation of trace indicates LED intensity. **l,** Input-output function curves of MEC_glu_ (blue) and LEC_glu_ (green) evoked CA3 PN EPSCs (−80 mV) and IPSCs (+10 mV) across all light intensities (IPSC MEC_glu_: n = 64 cells, N = 30 mice, LEC_glu_: n = 19 cells, N = 11 mice, NS p = 0.7605 Mixed-effects model w/ Geisser-Greenhouse Correction, EPSC MEC_glu_: n = 72 cells, N = 32 mice, LEC_glu_: n = 21 cells, N = 12 mice, NS p = 0.9629 Mixed-effects model w/ Geisser-Greenhouse Correction). **m,** Peak EPSC (−80 mV) and IPSC (+10 mV) amplitude of CA3 PN response to LEC_glu_ (green) and MEC_glu_ (blue) at low (8.8%) light intensity (left, EPSC: NS p = 0.3539 Mann-Whitney two- tailed, IPSC: NS p = 0.1898 Mann-Whitney two-tailed) and high (100%) light intensity (right, EPSC: NS p = 0.8492 Mann-Whitney two-tailed, IPSC NS p = 0.6865 Mann-Whitney two-tailed) to compare putative monosynaptic and polysynaptic pathway differences. Each data point represents the average PSC amplitude of an individual neuron. **n,** E/I ratio of MEC_glu_- (blue, n = 61 cells, N = 30 mice) and LEC_glu_-driven (green, n = 16 cells, N = 10 mice) CA3 PN EPSCs/IPSCs (−80 mV/+10 mV) at both low (8.8%, left, NS p = 0.8262 Mann-Whitney two-tailed) and high (100%, right, NS p = 0.7206 Mann-Whitney two-tailed) LED intensities. Each data point represents the average E/I ratio of an individual neuron and only neurons from which both EPSCs and IPSCs were recorded are included in the E/I ratio analysis. All error bars represent SEM.

To establish functional connectivity between the brain regions we optogenetically stimulated opsin-expressing MEC_glu_ and LEC_glu_ axons while performing whole-cell patch-clamp recordings from CA3 PNs in transverse entorhinal-hippocampal slices (Fig. 1c,e-h)^46,72^. All recordings were performed in CA3a/b PNs, and the borders of CA2 and CA3c were avoided. To stimulate Chronos and ChrimsonR expressing axons, we used 1ms-long light pulses at 470 nm and 625 nm wavelengths, respectively. We voltage-clamped CA3 neurons at −80 mV to isolate excitatory post-synaptic currents (EPSCs). We found that MEC_glu_ and LEC_glu_ stimulation elicited responses in 100% of CA3 PNs (n = 37 of 37 cells) in control (ACSF) conditions (Fig. 1e,f,h). To isolate monosynaptic connections, we bath applied TTX and 4AP, thereby blocking Na^+^ channels to eliminate polysynaptic responses and blocking K^+^ channels to allow opsin-mediated spike-independent transmission^73^. We found that all CA3 PNs except one (n = 20 of 21 cells) were monosynaptically connected to both LEC_glu_ and MEC_glu_ (Fig. 1f), confirming that the typical CA3 PN receives direct glutamatergic inputs from both MEC and LEC.

To determine whether MEC_glu_ or LEC_glu_ drives stronger excitation to CA3, we optically stimulated each input across a range of light intensities while recording EPSCs from CA3 PNs (Fig. 1g). Analysis was restricted to stimulation at low-light intensities to minimize opsin crosstalk, as blue light can activate ChrimsonR (red opsin) at higher intensities^7,46^ (Extended Data Fig. 1d-h). In pharmacological monosynaptic conditions (TTX + 4-AP), MEC_glu_ inputs evoked two times larger EPSC amplitudes (∼ −40pA) than LEC_glu_ (∼ −20pA, p = 0.0039). A similar trend was observed in ACSF, though it was not significant (Fig. 1g,h). This stronger MEC_glu_ drive onto CA3 is biophysically consistent with the more proximal termination of MEC_glu_ axons to the CA3 soma compared to LEC_glu_.

### MEC_glu_ and LEC_glu_ drive similar excitation and inhibition in CA3 PN

Next, we tested whether MEC_glu_ and LEC_glu_ recruit different amounts of excitation and feedforward inhibition (FFI) in CA3 PN using the previously described dual-optogenetic method (Fig. 2a). We interleaved opsins between pathways in the dual-labeled strategy to control for possible differences arising from opsin-specific effects. This dual-labelled method allowed us to look at responses to both inputs within an individual cell across a small range of light intensities. At low (<10%, 0.31mW/mm^2^) light intensities, we also avoided potential contamination via polysynaptic driven excitation^46^. To test the synaptic strength and excitation-inhibition ratio in these circuits, we voltage-clamped CA3 PNs at −80 mV and +10 mV, isolating EPSCs and inhibitory post-synaptic currents (IPSCs), and stimulated MEC_glu_ and LEC_glu_ axons with minimal stimulation (1 ms pulse) across a range (0-10%) of light intensities (Fig. 2b-h). We found that both MEC_glu_ and LEC_glu_ stimulation produced comparable EPSC amplitudes, confirming similar excitatory drive in the putative monosynaptic regime in ACSF. Additionally, we observed large light-evoked IPSCs with similar amplitudes driven by each pathway, suggesting recruitment of strong FFI onto CA3 PN (Fig. 2b,c). The excitation-inhibition ratio was <1 for both inputs and was not significantly different from each other (Fig. 2d). Thus, in ACSF conditions, permissive of polysynaptic activity, MEC_glu_ and LEC_glu_ drive similar levels of excitation and FFI in CA3 PNs, and the feedforward inhibition component dominates (Fig 2c,d).

To test whether the variability of MEC_glu_ and LEC_glu_ driven excitation-inhibition within an individual neuron reflects the population level similarities, we computed each neuron’s MEC-LEC response index (the ratio of the difference of the CA3 response to MEC_glu_ and LEC_glu_ to the sum of the CA3 response to MEC_glu_ and LEC_glu_). We plotted the EPSCs or IPSCs driven by MEC_glu_ as a function of the corresponding LEC_glu_ responses recorded from each neuron (Fig 2e,g). The average of all the cell response indices shows that MEC_glu_ drives more excitation and inhibition (response index > 0) at both −80 mV and +10 mV (stimulation intensities are <10%, Fig 2f,h). However, individually CA3 PN responses to MEC_glu_ are not categorically larger than LEC_glu_ indicating cell to cell variability within the CA3 PN population (Fig 2e,g).

Additionally, we analyzed the kinetics of the MEC_glu_- and LEC_glu_-driven responses and found that the two pathways are remarkably similar across all physiological properties we measured (Extended Data Fig. 2, Table 1,2). To avoid crosstalk and look at a larger range of stimulation intensities, we used a single-opsin strategy to label MEC_glu_ or LEC_glu_ in independent animals (single-labelled, Fig 2i-k). To compare MEC_glu_ and LEC_glu_ across animals, we virally expressed Chronos in glutamatergic neurons of either LEC or MEC and recorded CA3 PN responses to photostimulation of opsin-labeled MEC_glu_ or LEC_glu_ input. As before with dual-labelled experiments, MEC_glu_ and LEC_glu_ yielded similar PSC amplitudes at putative monosynaptic (low) light intensities. In addition, we observed the same results at polysynaptic (high) light intensities (Fig. 2l,m), something we were unable to test due to crosstalk in the dual-labelled experiments. This indicates that MEC_glu_ and LEC_glu_ to CA3 PN have similar excitatory and inhibitory drive at both direct and network levels across the population of recorded CA3 a/b PNs. Further, we confirmed that, as in the dual-optogenetic experiments, excitation-inhibition ratios were <1 and did not differ between LEC_glu_ and MEC_glu_, suggesting feedforward inhibition dominates in these two circuits at both the mono- and poly-synaptic levels (Fig. 2n).

**Table 1:** P-values for MEC_glu_- or LEC_glu_- driven CA3 response kinetics, input/output comparisons Extended Fig. 2.

|  | Labelling | Holding (mV) | p-value | Test | Fig. Panel |
| --- | --- | --- | --- | --- | --- |
| **Summation** | Dual | 80 | NS, 0.6434 | Mixed Effects Model<br>w/ Geisser<br>Greenhouse<br>Correction | b |
| **Summation** | Dual | 10 | NS, 0.5387 |  | b |
| **AUC** | Dual | 80 | NS, 0.9564 |  | c |
| **AUC** | Dual | 10 | NS, 0.7183 |  | c |
| **Decay** | Dual | 80 | NS 0.2337 |  | d |
| **Decay** | Dual | 10 | NS, 0.5594 |  | d |
| **Latency** | Dual | 80 | NS, 0.4645 |  | e |
| **Latency** | Dual | 10 | NS, 0.0987 |  | e |
| **Half-Width** | Dual | 80 | NS, 0.1738 |  | f |
| **Half-Width** | Dual | 10 | NS, 0.0788 |  | f |
| **Rise Time** | Dual | 80 | NS, 0.7329 |  | g |
| **Rise Time** | Dual | 10 | NS, 0.7004 |  | g |
| **Summation** | Single | 80 | NS, 0.7661 |  | l (left, middle) |
| **Summation** | Single | 10 | NS, 0.9605 |  | l (left) |
| **AUC** | Single | 80 | NS, 0.7505 |  | j (left, middle) |
| **AUC** | Single | 10 | NS, 0.6555 |  | j (left) |
| **Decay** | Single | 80 | NS, 0.2675 |  | k (left, middle) |
| **Decay** | Single | 10 | NS, 0.2478 |  | k (left) |
| **Latency** | Single | 80 | NS, 0.1091 |  | l (left, middle) |
| **Latency** | Single | 10 | NS, 0.7093 |  | l (left) |
| **Rise Time** | Single | 80 | NS, 0.8484 |  | m (left, middle) |
| **Rise Time** | Single | 10 | NS, 0.3932 |  | m (left) |
| **Half-Width** | Single | 80 | *, 0.0339 |  | n (left, middle) |
| **Half-Width** | Single | 10 | NS, 0.3555 |  | n (left) |

**Table 2:** Quantification and p-values of the MEC_glu_- or LEC_glu_- driven CA3 response kinetics, high/low comparisons Extended Fig. 2.

|  | Holding<br>(mv) | MEC<br>Mean<br>Value<br>(low) | LEC Mean<br>Value<br>(low) | MEC<br>Mean<br>Value<br>(high) | LEC Mean<br>Value<br>(high) | p-value<br>(low) | p-value<br>(high) | Test (low) | Test (high) | Fig. Panel |
| --- | --- | --- | --- | --- | --- | --- | --- | --- | --- | --- |
| **Summation** | 80 | -123.1 | -202.3 | -494.3 | -493.9 | NS, 0.1547 | NS, 0.9978 | Unpaired t test w/ Welch's Correction |  | N/A |
| **Summation** | 10 | 693 | 795.6 | 1058 | 994.1 | NS, 0.4070 | NS, 0.8466 | Mann-Whitney Test |  | N/A |
| **AUC** | 80 | -45.43 | -67.26 | -142.2 | -121.3 | NS, 0.3020 | NS, 0.4769 | Unpaired t test w/ Welch's Correction |  | N/A |
| **AUC** | 10 | 320.2 | 359.2 | 506.9 | 442.8 | NS, 0.6549 | NS, 0.6598 | Mann-Whitney Test |  | N/A |
| **Decay** | 80 | 16.63 | 21.04 | 25.55 | 26.43 | NS, 0.0668 | NS, 0.6488 | Unpaired t test w/ Welch's Correction |  | k (right) |
| **Decay** | 10 | 45.95 | 44.1 | 47.6 | 43.77 | NS, 0.2554 | NS, 0.0866 | Mann-Whitney Test |  | k (right, middle) |
| **Latency** | 80 | 10.36 | 7.697 | 6.241 | 5.184 | NS, 0.2289 | NS, 0.9288 | Lognormal Welch's t test |  | l (right) |
| **Latency** | 10 | 8.19 | 9.045 | 6.765 | 7.048 | NS, 0.2094 | NS, 0.8741 | Lognormal Welch's t test |  | l (right,middle) |
| **Rise Time** | 80 | 10.91 | 9.511 | 1.01 | 10.51 | NS, 0.7679 | NS, 0.8112 | Lognormal Welch's t<br>test | Mann-Whitney<br>Test | m (right) |
| **Rise Time** | 10 | 11.15 | 9.884 | 12.85 | 10.32 | NS, 0.1683 | **, 0.0050 | Unpaired t test with Welch's Correction |  | m (right,middle) |
| **Half-Width** | 80 | 11.5 | 15.43 | 18.76 | 20.85 | NS, 0.0727 | NS, 0.2029 | Mann-Whitney Test | Unpaired t test with<br>Welch's Correction | n (right) |
| **Half-Width** | 10 | 38.33 | 36.68 | 41.02 | 36.25 | NS, 0.5868 | **, 0.0080 | Unpaired t test with<br>Welch's Correction | Unpaired t test with<br>Welch's Correction | n (right, middle) |

### MEC_glu_ and LEC_glu_ have different frequency-dependent excitatory short-term plasticity

*In vivo*, MEC and LEC exhibit distinct patterns of task-relevant activity that are critical for communicating information across the cortico-hippocampal axis^49^. Therefore, we decided to explore the temporal dynamics of each input at the single-cell level. We asked whether MEC_glu_ and LEC_glu_ have different activity-dependent presynaptic properties measured by paired pulse ratios (PPR) with short-term plasticity (STP) induction. For this experiment, Chronos was expressed in each pathway individually due to its faster off-kinetics and higher frequency fidelity^7,74^. MEC_glu_ or LEC_glu_ axons in these single labeled animals were stimulated with trains of low-light intensity pulses (20 pulses, 1 ms duration, λ= 480 nm) at different frequencies (10 hz, 20 hz, and 40 hz) while recording CA3 PNs under voltage-clamp at both −80 mV and +10 mV (Fig. 3). We found MEC_glu_ and LEC_glu_ driven EPSCs both facilitated (PPR > 1) during the 10 hz train with increasing pulse number (P2-20) (Fig. 3a,b,d). However, at 20 hz, we see that while MEC_glu_ continues to facilitate across all pulses, LEC_glu_ starts to depress (PPR < 1) around pulse 5 (Fig. 3a,b,d). As we were unable to resolve time-locked peaks to pulses 3-20 in higher frequency trains in most cells, we compared the PPR of the first two pulses (P2/P1) across the different frequencies and the area under the curve (AUC) to measure overall synaptic drive.

**Figure 3:**
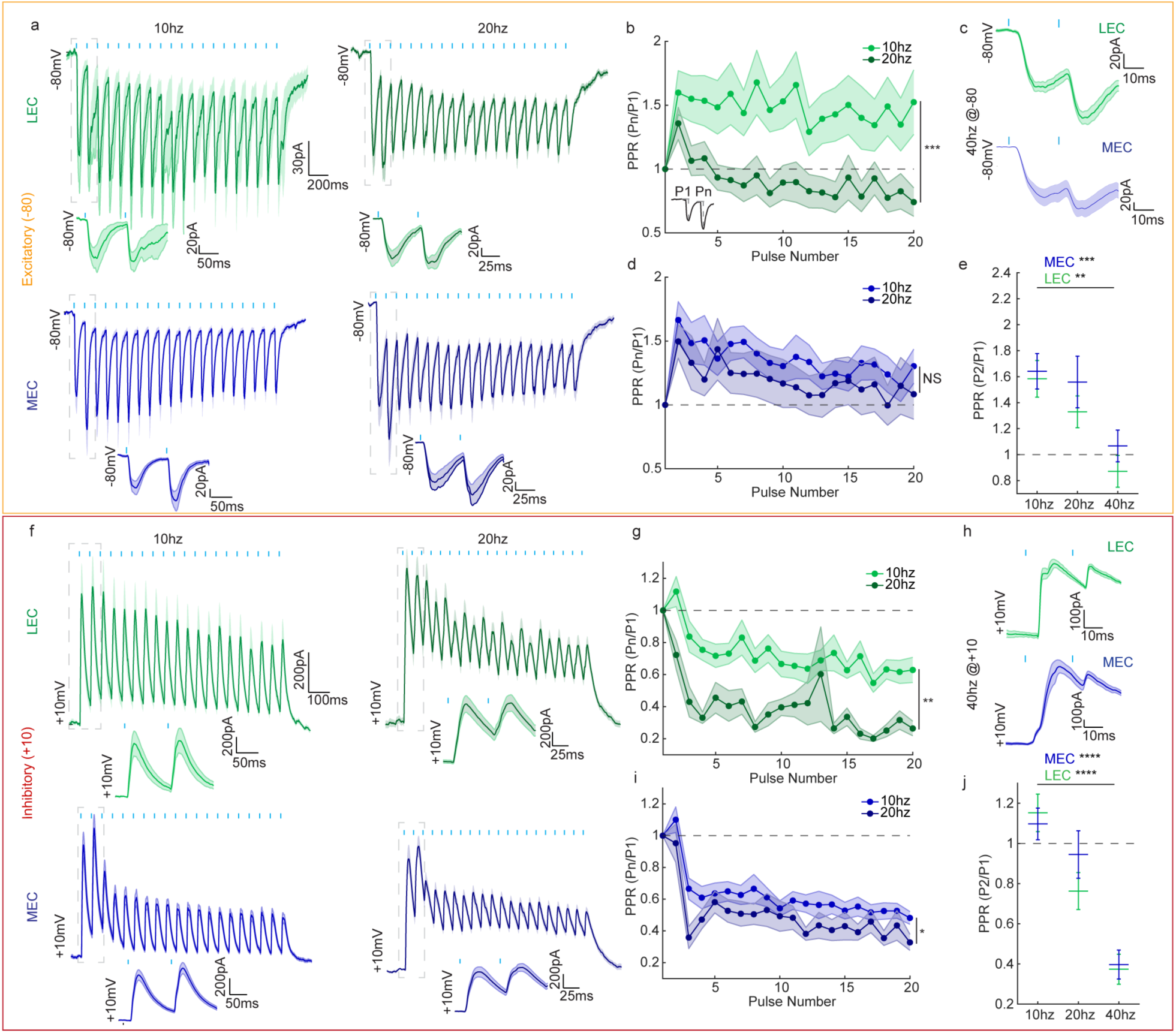
MEC_glu_ and LEC_glu_ have a frequency dependent divergence in short term excitatory plasticity. **a,f,** Averaged traces of CA3 PN responses to 10 hz (top) and 20 hz (bottom) LEC_glu_ (green) and MEC_glu_ (blue) stimulation in voltage-clamp at **a,** −80 mV and at **f,** +10 mV. Expanded view of demarcated area is responses to stimuli #1 and #2. **b,d** Summary plots of paired-pulse ratio (PPR) of the 1-20th MEC_glu_ (blue, **b**) and LEC_glu_ (green, **d**) driven EPSCs at 10 hz and 20 hz (MEC_glu_ 10 hz: n = 54 cells, N = 25 mice, 20 hz: n = 54 cells, N = 25 mice, *p = 0.0155 Mixed-effects model w/ Geisser-Greenhouse Correction; LEC_glu_ 10 hz: n = 23 cells, N = 13 mice, 20 hz: n = 20 cells, N = 12 mice, ***p = 0.0044 Mixed-effects model w/ Geisser-Greenhouse Correction). **g,i,** Summary plots of PPR of the 1-20th MEC_glu_ (blue, **g**) and LEC_glu_ (green, **i**) driven IPSCs at 10 hz and 20 hz (MEC_glu_ 10 hz: n = 35 cells, N = 19 mice, 20 hz: n = 32 cells, N = 18 mice, *p = 0.0477 Mixed-effects model w/ Geisser-Greenhouse Correction; LEC_glu_ 10 hz: n = 13 cells, N = 9 mice, 20 hz: n = 13 cells, N = 9 mice, **p = 0.0034 Mixed-effects model w/ Geisser-Greenhouse Correction). **c, h,** Sample averaged traces of CA3 PN excitatory responses to 40 hz stimulation of LEC_glu_ (green) and MEC_glu_ (blue) at **c,** −80 mV (**c**) and **h,** +10 mv. **e, j,** Paired-pulse ratio of the peak amplitude of **e,** 2nd EPSC and **j,** IPSC for MEC_glu_ (blue) and LEC_glu_ (green) at 10 hz, 20 hz, and 40 hz (MEC_glu_ EPSC 40 hz: n = 50 cells, N = 27 mice, ***p=0.0001 Kruskal-Wallis test; LEC_glu_ EPSC 40 hz: n = 15, N = 9 mice, **p=0.0025 one-way ANOVA; MEC_glu_ IPSC 40 hz: n = 26 cells, N = 18 mice, ****p < 0.0001 one-way ANOVA; LEC_glu_ IPSC 40 hz: 12 cells, N = 9 mice; ****p < 0.0001 one-way ANOVA). All error bars represent SEM.

With this early train metric, we found that MEC_glu_ and LEC_glu_ both facilitate at 10 hz and 20 hz, but at 40 hz LEC_glu_ depresses while MEC_glu_ facilitates even at higher frequencies (Fig. 3c,e). The total AUC of the responses to the train of stimulation for MEC and LEC grows with increasing frequencies for both the +10 mV and −80 mV conditions (Extended Data Fig. 3f,g). The ratio of the AUC between each stimulation pulse in the train compared to the AUC for pulse 1 stimulation at −80 mV is no different for MEC between 10, 20, and 40 hz, but we do see a difference in LEC in the 40 hz regime (Extended Data Fig. 3c). Thus, MEC_glu_ and LEC_glu_-excitatory synapses to CA3 PNs exhibit divergent short-term plasticity dynamics at higher frequencies.

We next examined the STP characteristics of MEC_glu-_ and LEC_glu_-driven FFI under the same stimulation paradigm (Fig. 3f-j). In contrast to −80 mV data, at +10 mV we found at 10 hz, MEC_glu_- and LEC_glu_-driven IPSCs initially facilitate (PPR>1 P_2_/P_1_) then depress (PPR<1, P_n_/P_1,_ n= pulses 3-20) with increasing pulse number (Fig. 3f,g,i). At 20 hz, we found that both MEC_glu_ and LEC_glu_ stimulation resulted in depression of IPSCs throughout the entire stimulation train (Fig. 3f,g,i). Thus, MEC_glu_ and LEC_glu_ driven FFI STP do not diverge from each other: they both become depressing at higher frequencies (20 hz and 40 hz, Fig. 3g-j). Therefore, MEC_glu_ and LEC_glu_ STP dynamics differ at the E-CA3 PN synapses but not the FFI-CA3 PN synapses. This could be due to differences in synaptic properties of MEC_glu_/LEC_glu_ to GABAergic (E-I) or GABAergic to CA3 PN (I-PN) and recruitment of overlapping GABAergic interneurons (IN)^75,76^.

### MEC_glu_ and LEC_glu_ recruit overlapping interneuron subtypes

Due to the observed similarities in FFI STP and the substantial FFI driven by both MEC_glu_ and LEC_glu_ we asked what GABAergic microcircuits modulate the MEC_glu_ and LEC_glu_ responses in CA3 PNs. Among the different IN subtypes in the hippocampus, we selected four major genetically defined GABAergic neuron subtypes in CA3 based on their relevance to cortico-hippocampal and somato-dendritic information processing^77-80^: parvalbumin (PV)^81^, cholecystokinin (CCK) ^82^, vasoactive intestinal peptide (VIP)^83,84^, and somatostatin (SST)^85,86^. We chose to chemogenetically manipulate these IN subtypes and read out their influence on MEC_glu_- and LEC_glu_-driven FFI onto CA3 PNs (Fig. 4a,b).

**Figure 4:**
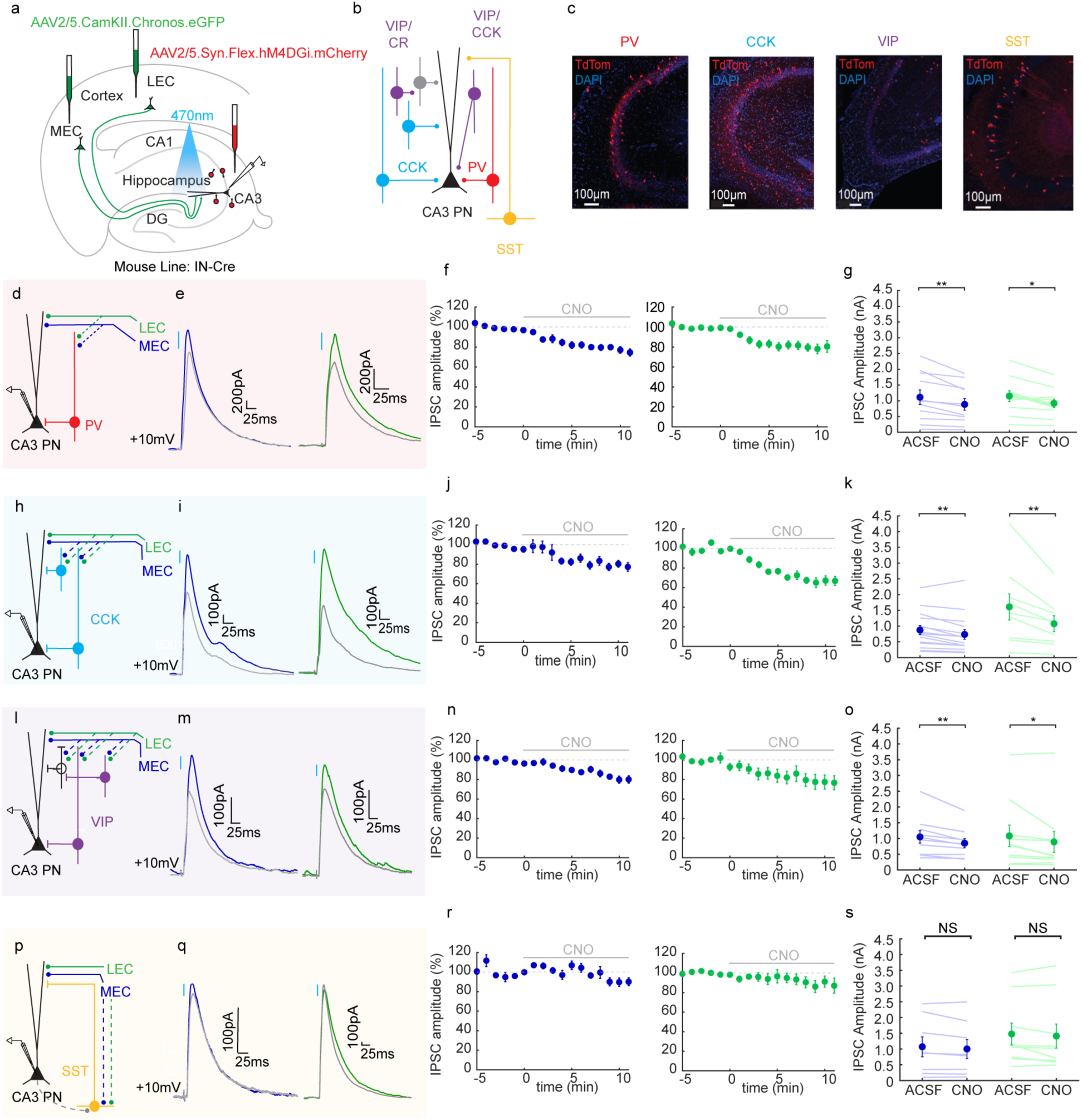
MEC_glu_ and LEC_glu_ to CA3 circuits are modulated by overlapping interneuron subtypes. **a,** Schematic of single-labelled viral injection strategy. AAV2/5.CaMKII.Chronos.eGFP was injected into MEC or LEC to drive excitatory opsin expression selectively in glutamatergic neurons. To label specific interneuron (IN) subtypes in CA3, a Cre-dependent Gi DREADD was injected in a GABAergic interneuron-Cre mouse line: parvalbumin (PV), cholecystokinin (CCK), vasoactive intestinal peptide (VIP), and somatostatin (SST). **b,** Circuit diagram of reported IN subtype connectivity with CA3 PN. **c,** Confocal images of CA3 region with representative distribution of interneuron subtypes labelled with tdTomato (red). PV INs are primarily in the *stratum pyramidale* (SP) region. CCK+ INs are found in SP, *stratum radiatum* (SR), and *stratum lacunosum moleculare* (SLM) of CA3. VIP+ INs in CA3 are sparse and mainly located in SP and SLM. SST INs are almost exclusively in *stratum oriens* (SO) and SP. **d-g,** PV+ IN silencing. **d**, Experimental schematic, putative circuits are dotted lines. **e**, Representative traces of MEC_glu_- (blue) and LEC_glu_- (green) driven CA3 PN IPSCs before and after (grey) PV+ IN silencing. **f,** Time course of MEC_glu_ (blue, n = 11 cells, N = 5 mice) or LEC_glu_ (green, n = 11 cells, N = 3 mice) evoked CA3 PN IPSC amplitudes with CNO (10 µM) application. **g,** Summary data of MEC_glu_- and LEC_glu_- driven CA3 IPSC amplitude before (ACSF) and after (CNO) PV+ IN silencing. Each data point represents the average response from an individual cell (MEC_glu_: n = 11 cells, N = 5 mice, **p = 0.0039 paired t-test; LEC_glu_: n = 11 cells, N = 3 mice, **p = 0.0027 paired t-test). **h-k,** Same as in **d** but with CCK+ IN silencing (MEC_glu_: n = 15 cells, N = 7 mice, **p = 0.0041 paired t-test; LEC_glu_: n = 9 cells, N = 5 mice, *p = 0.0129 paired t-test). **l-o S**ame as in **d-g,** but with VIP+ IN silencing (MEC_glu_: n = 10 cells, N = 5 mice, **p = 0.0039 Wilcoxon matched-pairs two-tailed; LEC_glu_: n = 10 cells, N = 5 mice, *p = 0.0273 Wilcoxon matched-pairs two-tailed). **p-s, Same** as in **d-g,** but with SST+ IN silencing. (MEC_glu_: n = 8 cells, N = 6 mice, NS p = 0.0977 Wilcoxon matched-pairs two-tailed; LEC_glu_: n = 9 cells, N = 4 mice, NS p = 0.3594 Wilcoxon matched-pairs two-tailed). All error bars represent SEM.

To chemogenetically silence specific IN populations, we virally expressed a Cre-dependent G-protein coupled inhibitory designer receptor exclusively activated by designer drugs^87^ (Gi-DREADD) in CA3 of PV^+^, CCK^+^, VIP^+^, and SST^+^ Cre-driver mouse lines (Fig. 4c). In the same animals, we expressed Chronos in MEC_glu_ or LEC_glu_ (single-labelled) to optogenetically activate one input or the other (Fig. 4a,d,h,l,p). We recorded the CA3 PN responses to photo-stimulation of individual inputs at +10 mV to record light evoked IPSCs in the absence of (control) or presence of (silenced) the synthetic Gi-DREADD ligand (CNO, 10µM) via bath application (Fig. 4e,i,m,q). This allowed us to assess the contribution of each IN subtype to the MEC_glu_- or LEC_glu_-driven FFI within the CA3 circuits (Fig 4. f,j,n,r). Thus, we were able to record the response to MEC_glu_ or LEC_glu_ with and without specific CA3 interneuron subtypes intact.

We observed a reduction in MEC_glu_- and LEC_glu_-driven IPSC amplitude with PV^+^, CCK^+^, and VIP^+^ IN silencing (Fig 4d-o). In contrast, we did not observe a significant reduction in IPSC amplitude with SST^+^ IN silencing (Fig. 4p-s). We concluded that MEC_glu_-CA3 and LEC_glu_-CA3 circuits are both modulated by PV^+^, CCK^+^, and VIP^+^ IN subtypes. Unlike the other subtypes, SST^+^ INs were not recruited by the MEC_glu_ nor LEC_glu_ to CA3 circuit with our stimulation paradigm (Fig. 4p-s). Therefore, MEC_glu_ and LEC_glu_ likely converge on to the same GABAergic subtypes, consistent with their similar inhibitory temporal dynamics (STP) and FFI drive on to CA3 PNs.

### MEC_glu_ and LEC_glu_ recruit similar subthreshold output but rarely drive CA3 PN output

While MEC_glu_ excitatory drive on to CA3 PN is larger than LEC_glu_ in a pharmacologically-isolated monosynaptic regime (TTX & 4-AP), both inputs drive similar amounts of excitation and FFI in CA3 PN in regular ACSF. Moreover, these inputs exhibit different temporal dynamics (STP) at the direct EC-CA3 PN synapse, making it difficult to predict what the net CA3 PN output may be. We examined the influence of MEC_glu_ and LEC_glu_-driven excitation and inhibition on CA3 PN output. We used our dual-labelled strategy as previously described and performed whole-cell current-clamp recordings, adjusting to maintain membrane potential near −70 mV to control for driving force (Fig. 5a-c). Once again, low light intensities (λ =470 nm and 625 nm) were used to ensure minimal contamination from opsin crosstalk and polysynaptic excitation. The MEC_glu_ and LEC_glu_ driven sub-threshold responses were not significantly different across a range of low light intensities (Fig. 5c).

**Figure 5:**
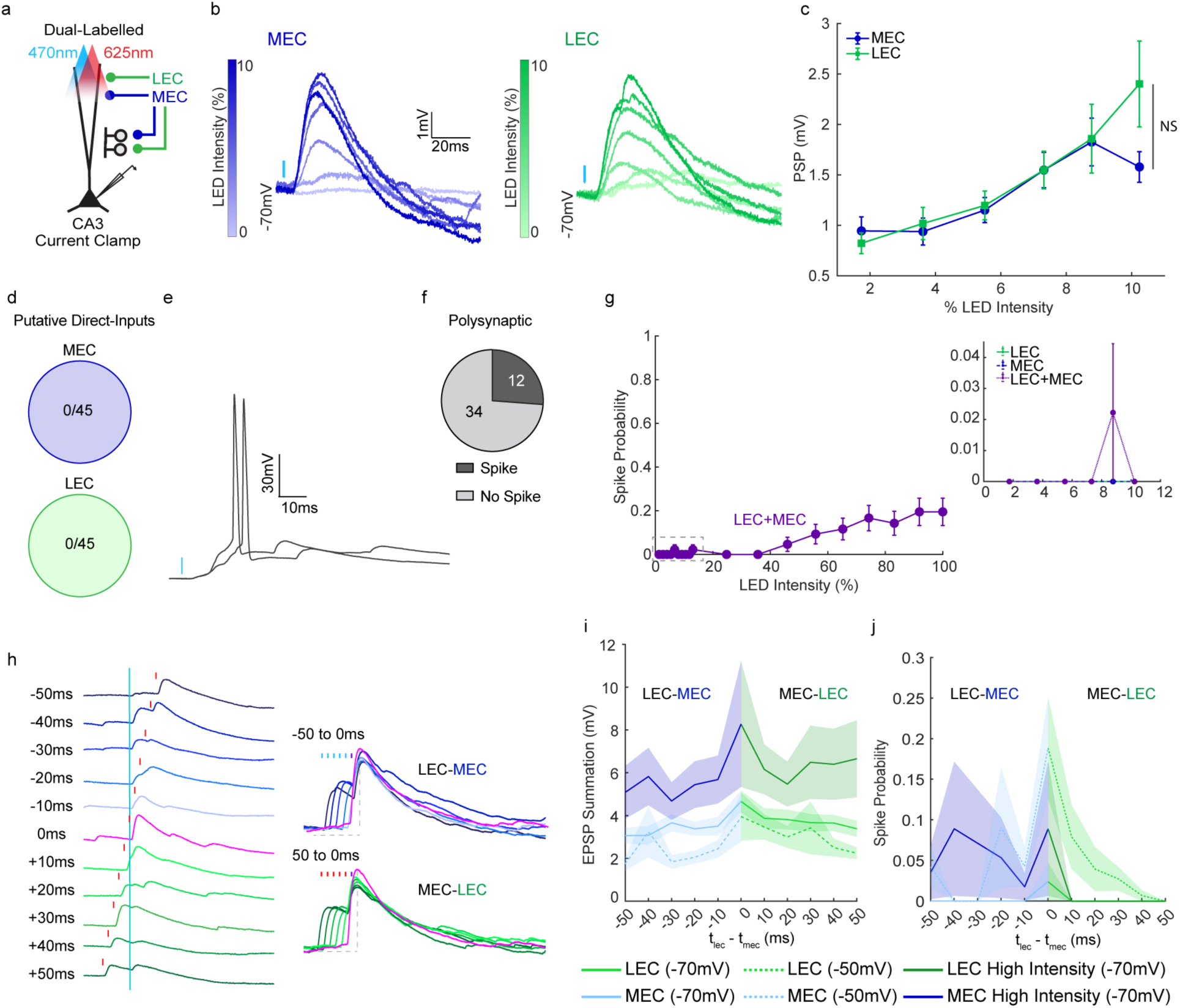
MEC_glu_ and LEC_glu_ putative monosynaptic connections do not drive action potentials in CA3 PN and the optimal window for integration is simultaneous. **a,** Dual-labelled experimental schematic. We recorded responses of CA3PN current-clamped at −70 mV to MEC_glu_ (blue) and LEC_glu_ (green) stimulation using 470 nm and 625 nm light over the axons. **b,** Representative traces of MEC_glu_- and LEC_glu_- driven CA3 PN PSPs at −70 mV across a range of low (1-10%, 0.05-0.31 mW/mm^2^) light intensities. Responses to MEC_glu_ and LEC_glu_ are from the same cell/animal. **c,** Input-output function curves of MEC_glu_ (blue) and LEC_glu_ (green) light-evoked CA3 PN PSPs across low light intensities. Only low light intensities are shown to prevent opsin crosstalk contamination (n = 45 cells, N = 21 mice; NS p= 0.5153, Mixed Effects Model w/ Geisser- Greenhouse Correction). **d,** Fraction of CA3 PNs that produced action potentials from MEC_glu_- or LEC_glu_- low light stimulation (n = 45 cells, N = 21 mice). **e,** Representative action potentials from CA3 PN upon coincident stimulation of both inputs (MEC_glu_ + LEC_glu_). **f,** Fraction of CA3 PNs that generate action potentials from MEC_glu_-, LEC_glu_-, or a combination stimulation at polysynaptic/high (10-100%, 0.31-3.02 mW/mm^2^) light intensities (n = 46 cells, N = 21 mice). **g**, Input-output function curves of CA3 PN action potential firing probability in response to MEC_glu_ (blue), LEC_glu_ (green), and combined MEC_glu_ + LEC_glu_ (purple, 470 nm + 625 nm) stimulations across low light intensities (inset) and combined MEC_glu_ + LEC_glu_ stimulations at high light intensities. **h,** Representative traces of CA3 PN PSPs evoked from stimulation of MEC_glu_ and LEC_glu_ with interstimulus intervals (ISIs) of 0, 10, 20, 30, 40, and 50 ms. Baseline Vm was held at −70 mV. **i,** CA3 PN PSP summation amplitude as a function of ISI. **j,** Spike probability as a function of ISI. **i,j,** CA3 PN were recorded at holding potentials of −50 mV (dotted, light color) and −70 mV (solid lines). At −70 mV, both low (light color) and high intensity (dark color) LED stimulations were used (−70 mV low: n = 21 cells, N = 11 mice; −70 mV high: n = 7, N = 6 mice; −50 mV low: n = 13 cells, N = 10 mice). At high light intensities, it is not possible to attribute the response to only LEC_glu_ or MEC_glu_ due to opsin cross talk. All error bars represent SEM.

We then asked if the MEC and LEC inputs to CA3 were facilitating or depressing when both excitation and inhibition were at play. When measuring the paired-pulse ratio (PPR) at the highest light intensity below crosstalk at 10 hz, we found that both MEC_glu_ and LEC_glu_ to CA3 were facilitating on average (PPR > 1, Extended Data Fig. 4), consistent with the direct EC-CA3 PN paired-pulse facilitation seen in voltage-clamp (Fig. 3b,d). Furthermore, we examined the summation ratio to assess the post-synaptic depolarization driven by MEC_glu_ and LECglu in CA3 PNs and found that both inputs resulted in a depolarization buildup, with a summation ratio > 1 (Extended Data Fig. 4).

We then asked which of the two, MEC_glu_ or LEC_glu_, is more effective in driving spike output in CA3 PN. We performed spike detection on our low-intensity data to look at suprathreshold activity under putative monosynaptic conditions. We found that neither MEC_glu_ nor LEC_glu_ yielded action potentials in CA3 PNs (n=46 cells, Fig. 5d). To simulate ongoing network activity, we tested spike output in polysynaptic regimes by stimulating both inputs independently and in combination (λ = 470 nm + 625 nm) using high light intensities. This approach likely recruits the MEC_glu_ and LEC_glu_ to DG circuit as well as recurrent activity in the CA3 PN network and cannot guarantee that we are only stimulating MEC_glu_ or LEC_glu_ when using one wavelength of light. We found that in ∼26% of cells (n=12/46 cells) MEC_glu_, LEC_glu_, or a combination of both inputs evoked action potentials in CA3 PNs (Fig. 5e-g). This implies that the MEC_glu_ and LEC_glu_ to CA3 circuit requires synchronous drive from multiple inputs to drive output (i.e. network activity) and is supported by our findings that MEC_glu_ and LEC_glu_ inputs drive large amounts of FFI in CA3 PN.

### MEC_glu_ and LEC_glu_ have a narrow optimal time window for integration

Since integration across multiple synapses appears to be necessary for MEC_glu_ and LEC_glu_ to elicit CA3 PN action potentials, we explored the temporal window of integration for these two inputs by looking at the effect of convergent activity of MEC_glu_ and LEC_glu_ on CA3 PN depolarization and spike output. We used dual-labelled animals and stimulated each input with varying interstimulus intervals (ISIs) between the two inputs (0-50 ms) while recording from CA3 PN in current-clamp at −70 mV (Fig. 5h). As ISIs became shorter, the summation became larger, with the peak depolarization at 0 ms (Fig. 5i). Since this temporal summation window was very narrow, we asked if holding the cell’s membrane at more depolarized (−50 mV) *in vivo* like potentials would boost summation to better reach spike threshold with broader temporal integration. Such depolarization did not result in a change in the temporal window for integration nor did it facilitate an increase in summation of net subthreshold depolarization as compared to the −70 mV condition (Fig. 5i). However, the depolarized CA3 PN did show an increased spike probability (18.9%) and broadening of the timing window for integration to elicit spike output (Fig. 5j). Finally, we performed this experiment with a high light intensity stimulation at −70 mV to better approximate a polysynaptic regime where stimulations may recruit the EC to DG circuit and the CA3 PN recurrent network in addition to the direct EC-CA3 PN inputs. The CA3 PN responses in this paradigm are not exclusive to the individual input pathways due to opsin crosstalk (Fig. 5i,j, Extended Data Fig. 1). Recruitment of polysynaptic circuits did boost the summation but did not influence the temporal window for integration as compared to the low-intensity conditions (Fig. 5j). Similar to the depolarized condition, we found an increase in spike probability (8.9%) with high light intensity stimulation when MEC_glu_ and LEC_glu_ are stimulated simultaneously, ISI = 0 ms (Fig. 5j). Consistent with single-cell integration of distal innervation, modest excitation, and substantial inhibition, network effects are needed for MEC_glu_ and LEC_glu_ to spike CA3 PNs.

### *In vivo* the CA3 response to MEC-originating input is greater than the response to LEC-originating input

Despite displaying different temporal dynamics, MEC_glu_ and LEC_glu_ direct inputs to CA3 mostly share similar properties and rarely drive spike output *ex vivo*. Given that they convey different sensory modalities, this stalemate motivated us to examine their dynamics within an intact network *in vivo* where the entorhinal-hippocampal circuit has ongoing activity that is organized by prominent oscillations in the local field potential (LFP)^88^. Are MEC- and LEC-originating inputs to CA3 timed preferentially or independently with respect to ongoing fluctuations in network excitability (Fig. 6a)? We took advantage of a published dataset^48^ in which current source density source-localization identified dentate spikes (DS) as spontaneous synchronous discharge events that originate in either LEC (DS_LEC_) or MEC (DS_MEC_; Fig. 6b)^48,89^. LEC and MEC layer II inputs target both DG and CA3 areas, but are naturally segregated in DG. This allowed us to investigate the population discharge response of CA3 cells to DS_LEC_ and DS_MEC_ as a proxy for LEC and MEC activation of CA3 and their sub-second combinations in the awake intact brain. The CA3 response time-locked to DS_LEC_ events was weaker compared to the response time-locked to DS_MEC_ events, whether we compared the probability of spiking (max) or min/max difference in spike probability within a small window centered on the peak response time (−4 ± 20 ms; Fig. 6c, d). There was theta-range (∼140 ms) modulation of spike probability around each DS event with DS_MEC_ appearing near the peak of the modulation, and DS_LEC_ occurring in the trough, which may underlie their differential driving of CA3 firing.

**Figure 6:**
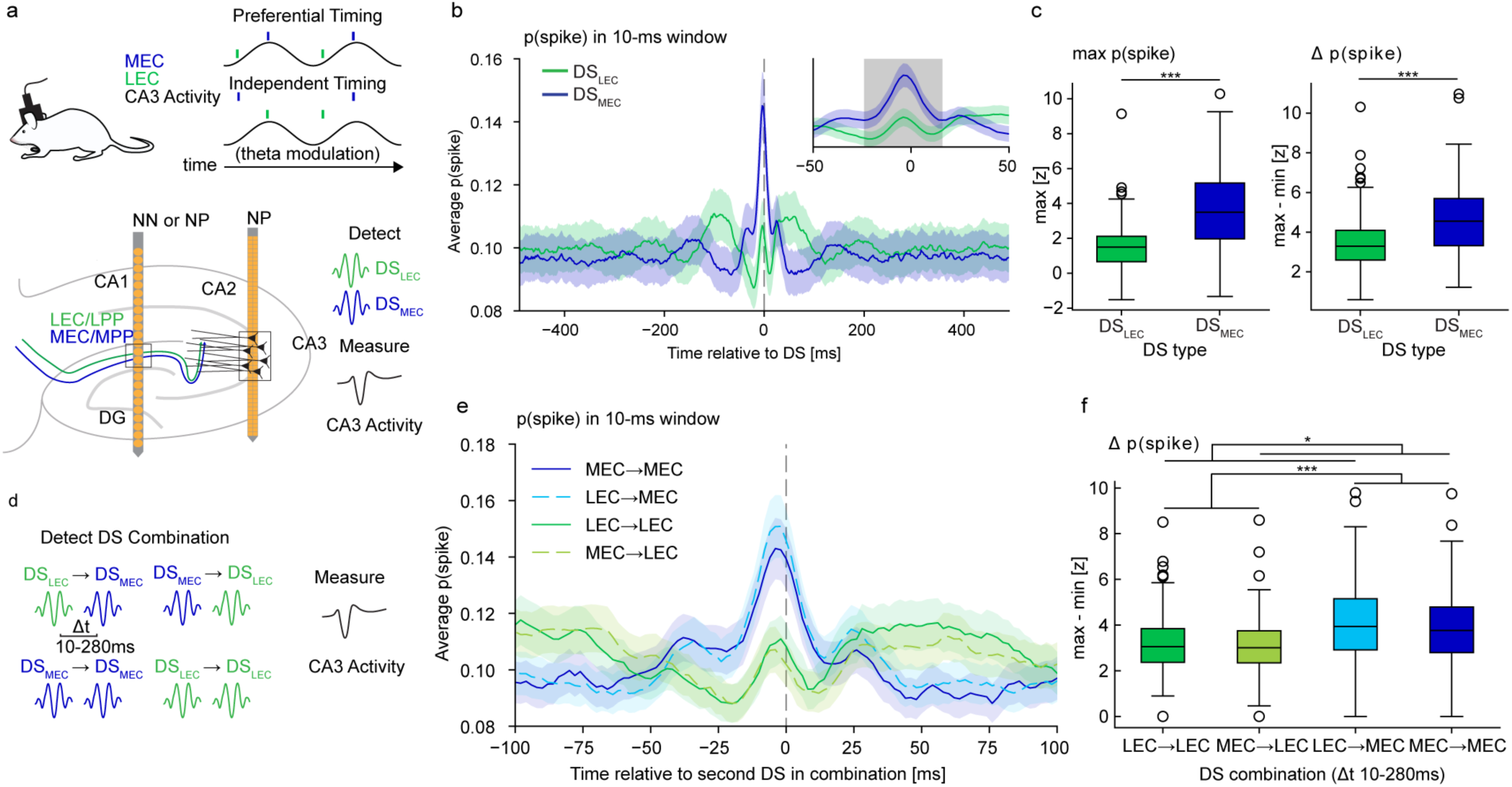
Greater CA3 responses to spontaneous MEC- versus LEC-originating dentate spikes in awake mice. **a,** Schematic of preferential (top) or independent (bottom) phase-locking of MEC (blue) and LEC (green) to CA3 theta-modulated activity (black). (bottom) Schematic of in-vivo electrophysiology recordings and analysis: Neuronexus linear electrode array (NN) or Neuropixels (NP) placement in CA1/DG for detection of DS events, NP placement in CA3 for measuring CA3 activity. Dentate spikes originating in MEC or LEC were detected then the CA3 activity in that time window was analyzed. **b,** Peri-DS time histogram of CA3 action potential probability from DS events originating in LEC (green, DS_LEC_) or MEC (blue, DS_MEC_). p(spike) at time t is the probability of observing one or more action potentials within a 10 ms window centered on t (step size 2 ms). Error bars: standard error metric (n=257 cells from 3 mice). **c,** Response strength was quantified for each cell by z-scoring its p(spike) trace over a 1- second interval centered on t=0 and then calculating either the max (left) or difference between max and min (right) observed over a 40-ms wide window centered on the time point of maximal population response (−4 ± 20 ms; gray shaded region in b). DS_MEC_ yields larger responses than DS_LEC_ (left: ****p = 1.5 × 10-22, right: ****p = 2.6 × 10-17 Wilcoxon signed-rank test,). **d,** Schematic of DS_MEC/LEC_ combinations that were detected with corresponding CA3 activity measured and quantified in e, f. **e,** Peri-DS time histogram of CA3 action potential probability from the second DS (“current”) in each possible combination of consecutive DS events, i.e. DS_LEC_ (green) or DS_MEC_ (blue) preceded by the same (dark) or different (light) type of DS (“preceding”). Error bars: standard error metric (n=257 cells). **f,** Response strengths were quantified as described in c (right), i.e. using the difference between max and min z-scored p(spike) over the window t = −4 ± 20 ms. A two-way ANOVA was used to evaluate differences in response strength given the “preceding” (i.e. first in combination) DS type, “current” (i.e. second in combination) DS type, and interaction between these two categorical variables (levels: DS_LEC_, DS_MEC_ for both variables). Current DS: F1,256 = 86.856, ***p = 0.00; preceding DS: F1,256 = 5.071, *p = 0.0252; interaction: F1,256 = 0.559, NS p = 0.455. Extended Fig. 5e shows the same analysis with response quantified using max z-scored p(spike), as described in c (left).

To parse out interactions between LEC and MEC drives *in vivo*, we examined these CA3 spiking probabilities in relation to preceding DS events (Fig. 6e). Whether we examined combinations of DS events that appeared within 10-280 ms (2 theta cycles; Fig. 6f, Extended Data Fig. 5e) of each other or 10-50 ms (Extended Data Fig. 5a,b), we came to the same conclusion. As before, DS_MEC_ drove more CA3 firing than DS_LEC_, regardless of whether they are preceded by DS_MEC_ or DS_LEC_ events. Further, the CA3 responses to either DS events following a DS_MEC_ were smaller than those following a DS_LEC_, indicating hysteresis due to preceding DS_MEC_- induced firing. In addition, the response to preceding DS_MEC_ events is evident at 50 ms before either DS event (Fig. 6Sa). We examined the timing (Extended Data Fig. 5c) and likelihood (Extended Data Fig. 5d) of DS event combinations. DS_LEC_ events (59.61%) are more common than DS_MEC_ events (40.39%), and the likelihood of their combinations are not consistent with pairs of events being independent. The histogram of inter-DS intervals is log-normal and bimodally distributed, related to theta cycle duration such that different DS events are more likely to appear within the same theta cycle than the same type of DS event (Extended Data Fig. 5c). Overall, we find that in freely behaving mice, the CA3 network responds more strongly to MEC activation than LEC activation.

## Discussion

Here we investigated how MEC and LEC are processed in hippocampal area CA3, whether they are functionally integrated or their segregation is maintained, and their consequences on network activity. Using dual-colored optogenetics and *ex vivo* electrophysiology, we show that CA3 PNs receive monosynaptic connections from both MEC and LEC glutamatergic inputs.

Underscoring that integration of multimodal inputs is relevant at a single cell level. We found that the wiring and output of these circuits are largely similar at the single neuron level, with two exceptions. First, consistent with the dendritic loci of their synapses onto CA3 PNs, the pharmacologically-isolated, monosynaptic, excitatory drive of MEC is slightly stronger than that of LEC. Second, MEC and LEC inputs to CA3 PNs exhibit different activity-dependent temporal dynamics: LEC_glu_ synapses facilitate at 10 hz but switch to a depression at 20 hz with increasing stimulation number, whereas MEC_glu_ synapses continue to facilitate at the higher frequency. Despite the similarities in synaptic wiring and biophysical properties of these two inputs observed *ex vivo* at the single-cell level, we found that *in vivo* CA3 suprathreshold network activity is more strongly driven by MEC-originating dentate spikes (DS_MEC_) than by LEC-originating dentate spikes (DS_LEC_). In contrast to DS_LEC_, DS_MEC_ events reliably evoked CA3 action potential discharge. DS_MEC_-associated activity was greater irrespective of the order and interval between LEC and MEC-originating dentate spikes. Our findings indicate that the hard- wiring of MEC_glu_ and LEC_glu_ to CA3 is largely similar, but the temporal details of these two inputs may enable them to relay their input-specific sensory information.

### MEC and LEC circuits in the hippocampus

A central dogma in the field is that information originating in MEC and LEC systems maintain distinct anatomical and functional organization within the hippocampus^8,13,56,59,90^. Our study investigated whether the two pathways remain distinct in CA3 using dual color optogenetics, allowing for within animal and cell comparisons. In terms of circuit design, the hippocampus could implement MEC/LEC input integration in three conceptually distinct ways: i) differential wiring to subsets of neurons which recurrently connect within CA3, ii) combining the information downstream via the CA3-CA1 trisynaptic circuit, or iii) by converging at the level of the single neuron (i.e. differential wiring to subsets of synapses within the same cell). Our results were unambiguous: both pathways target the same CA3 neurons, therefore integration of qualitatively different sensory information happens at the level of single principal neurons in CA3 rather than downstream at the network level. This implies that these two pathways likely do not recruit separate CA3 PN ensembles.

Our findings also showed that MEC drives more excitation in a pharmacologically-isolated, monosynaptic (TTX + 4-AP) regime, but in regular (ACSF) conditions, MEC_glu_ and LEC_glu_ excitatory and inhibitory drive onto CA3 PN is similar. This is consistent with decreased dendritic attenuation in TTX + 4-AP conditions yielding a more electrically compact membrane, thereby simply unveiling dendritic distance dependent-differences^91-94^. Further, this validates the sensitivity of our approach to detect minute differences despite the low resistivity and large volume of CA3 PNs. We note there is variability among individual CA3 PN responses to LEC_glu_ and MEC_glu_ stimulation. This could stem from the anatomical location of the CA3 PN (deep versus superficial)^95^ and/or the presence or absence of thorny excrescences^96-98^, which further studies could address.

In contrast to CA3, LEC was shown to drive greater activity in CA2 PNs than MEC using single pathway optogenetics^55^. While this approach did not allow for direct comparison of both pathways within a single neuron, this difference in circuit output may reflect functional distinctions between CA3 and CA2^99^. As CA2 is important for social memory behaviors, the contextual input (LEC) that provides odor information may be more relevant to CA2 than it is to CA3^33-35,70,100^. Perhaps CA3 relies more on the association of multimodal inputs from MEC and LEC due to its role in spatial episodic memory and context-dependent place cell _coding46,50,52,54,95,101._

Overall, our findings support the idea that both MEC_glu_ and LEC_glu_ behave as convergent skip connections to the vast majority of individual CA3 PN, allowing incoming cortical sensory information to be matched with that routed through DG via the mossy fiber inputs to provide a teaching signal^66^. Beyond the utility of a teaching signal, the stabilization of representations in CA3 by a skip connection would be especially valuable given the hypothesized lossy information processes of neurogenesis, pattern separation, sparsification, and regularization in DG^102-107^. Additionally, a complete copy of sensory information at the level of every CA3 PN, such that those selected by DG input are privy to both spatial and contextual information, could provide further specificity in ensemble selection before amplification via the CA3 recurrent collaterals.

### MEC_glu_ and LEC_glu_ to CA3 are modulated by overlapping GABAergic microcircuitry

Though it has been shown that different long-range inputs can preferentially target specific GABAergic neuron subtypes^108^, we found that MEC_glu_ and LEC_glu_ inputs recruit overlapping GABAergic interneuron subtypes. However, these parallel circuits (PV+, CCK+, and VIP+) could differentially modulate CA3 PN excitability in a context- and state-dependent manner according to the molecular makeup of each subtype^109,110^. PV+ INs are known to be fast-spiking and capable of high-frequency firing^81^. Though our data shows they modulate both the MEC_glu_ and LEC_glu_ to CA3 circuits, PV+ INs are one of the only subtypes that has the potential to keep up with the known *in vivo* gamma frequency of MEC (>100 hz)^49^. Further studies should address whether they preferentially follow MEC_glu_ inputs at higher frequencies in CA3.

In contrast, CCK-expressing GABAergic neurons have a slower firing frequency, are sensitive to endocannabinoids showing strong suppression of GABA release when CB1 receptors are activated, and their asynchronous GABA release prevents them from enforcing temporal fidelity in their transmission^111-115^ during repetitive stimulation. Thus, despite their ability to strongly modulate basal somatic and dendritic excitation^38,82,83,116^, at high frequencies CCK+ interneurons are likely offline and not benefiting from the increased facilitation of MEC_glu_ inputs unlike PV+ interneurons which can keep up with high frequency stimulation. Additional studies have shown CCK+ GABAergic neurons to be important for controlling theta phase precession and dampening sharp wave ripple power, but are thought not to be involved in gamma oscillatory changes^116-119^. Whereas PV+ cells are critical to maintaining both theta and gamma rhythms^120-122^. Because the MEC_glu_ and LEC_glu_ to CA3 circuit is modulated by both subtypes, these direct inputs likely have a bidirectional role in regulating temporal coding across oscillatory states in CA3.

Vasoactive Intestinal Peptide (VIP) INs are known for their inhibitory and disinhibitory role in the hippocampus^80,84,123,124^. In CA1 and CA3, LEC preferentially targets specific subtypes of VIP INs which allows for bidirectional modulation of hippocampal PN excitability^46,83^. While we did not dissect the subtype specificity of VIP INs, further studies could investigate whether MEC_glu_ and LEC_glu_ targeting of VIP subtypes diverge.

Studies have shown SST+ INs are recruited via feedback excitation from PNs^125-127^. As we did not observe any MEC_glu_- or LEC_glu_-driven CA3 action potentials at low intensities and only found that <30% of cells spike to high intensity stimulation, it is unsurprising we found that SST⁺ GABAergic neurons are not directly modulated by these circuits. Additionally, this is consistent with previous work from our lab showing that SST+ neurons in CA1 do not receive direct inputs from LEC_glu_ ^83^.

### Frequency dependent divergence in STP of MEC_glu_ and LEC_glu_ synapses in CA3

During trains of repetitive stimulation at varying frequencies, MEC_glu_ and LEC_glu_ synaptic responses show a frequency-dependent shift in their paired-pulse ratio. MEC_glu_ input to CA3 facilitates at 10, 20, and 40 hz with increasing pulse number, while LEC_glu_ input to CA3 facilitates at 10 hz, but this switches to depression at higher frequencies (20 and 40 hz) with increasing pulse number. On the other hand, STP properties of MEC_glu_- and LEC_glu_-driven FFI are remarkably similar to each other: facilitating at 10 hz but depressing at 20 and 40 hz. This could be an outcome of the overlapping GABAergic neurons that both pathways recruit within CA3. At higher frequencies the underlying suppression of FFI due to short term depression of inhibitory drive in combination with the excitatory facilitation of MEC_glu_ could provide a temporal window for continued excitation of the network via MEC_glu_ inputs as seen in the polysynaptic *in vivo* data.

Overall, our results suggest that CA3 neurons may rely on temporal codes to distinguish between the MEC_glu_ and LEC_glu_ inputs. This divergence in STP dynamics could reflect differences in the presynaptic molecular machinery. Some candidate molecules that allow for coupling of the MEC_glu_ and LEC_glu_ synaptic boutons with CA3 PN spines are: Munc 13-1 or Munc 13-2^128-130^, Synaptotagmin 1 vs Synaptotagmin 7^131^, or VGlut1 vs VGlut2^62,132^ isoforms. These are known to bestow distinct vesicular release probability, and synchronicity to synaptic terminals which could underlie the differences we observe in their STP. In addition, differences in the ratio of post-synaptic AMPA and NMDA receptor expression and activation^133^, in conjunction with recruitment of metabotropic Gq pathways and calcium-induced calcium release from intracellular stores,^82^ may also shape temporal and integrative dynamics of the distal dendritic MEC_glu_ and LEC_glu_ inputs during periods of sustained activity at higher frequencies.

Our observed behavior of LEC_glu_ input to CA3 is consistent with reports in DG, where Lateral Perforant Path-DG synapses act as a low pass filter such that responses to gamma frequencies are progressively suppressed^62^. Persistent facilitation at high frequencies may allow MEC_glu_ to continue to transfer information even during burst firing and high frequency PN activity^131^. Differences in paired pulse facilitation and neuromodulation between MEC and LEC synapses have also been reported in CA1^134^. Additionally, MEC and LEC inputs show distinct gamma frequency coupling *in vivo* with both CA1 and DG neurons in a task dependent manner. Fast gamma oscillations synchronize MEC with proximal CA1^69,135^ and DG granule cells^49^, whereas slow gamma oscillations coordinate LEC activity with CA1 during odor-place association learning ^70^ and DG mossy cells as well as CA3 PNs in an object-oriented learning task ^49^. Future studies are needed to define the mechanisms for filtering and/or integrating signals from MEC and LEC and test their consequences on ensemble coding and behavioral output.

### Linking the functional impact of MEC and LEC at cellular, circuit, and network levels

Our analyses of CA3 responses to spontaneous dentate spikes in the awake behaving mouse shows that MEC-originating dentate spikes are better at driving hippocampal output than LEC-originating dentate spikes^48^. We extended this relationship by examining the responses to DS_MEC_ and DS_LEC_ at various theta-related time intervals. We found that CA3 output is greater following DS_MEC,_ no matter the time interval between the DS_MEC_ and DS_LEC_. Consistent with the notion of independent responses to these two inputs, our corresponding *ex vivo* experiments show that the best window for MEC_glu_ and LEC_glu_ integration is when they arrive coincidently, a condition we could not reliably identify in the awake mouse data set due to the process of identifying dentate spike events. In this context, it is interesting to note that a third type of dentate spike has been identified where the inputs from MEC and LEC are coincident^89^. Our *ex vivo* work shows a small difference in direct input strength of MEC_glu_ over LEC_glu_, reflective of the anatomy. It is possible that this difference is further amplified by the CA3 recurrent circuitry and synaptic input barrages *in vivo*, where polysynaptic network-level activity motifs are engaged selectively by ongoing neuromodulatory and behavioral states. However, we cannot, beyond speculation, directly link our *ex vivo* within-cell observation that anatomically proximal MEC inputs drive stronger monosynaptic excitation than LEC, to our *in vivo* findings, that MEC drives more CA3 network activity than LEC. Our *ex vivo* findings rule out circuit design and differential recruitment of E/I balance by MEC_glu_ and LEC_glu_ to account for the dominance of MEC_glu_ over LEC_glu_ in generating CA3 responses *in vivo*, pointing to network effects.

One possible contribution is the emergent differences in short-term plasticity at the single-cell level where MEC_glu_-driven output is amplified at higher frequencies, but LEC_glu_-driven output is not. An additional possibility implicated by our data is that DS_MEC_ events are associated with an increase in CA3 spike probability, as compared to DS_LEC_ events, due to their timing relative to theta-modulated CA3 excitability. This could mean that, at a network level, the timing of inputs relative to the global phase of the network’s excitation cycle determines its ability to discriminate and integrate information. It seems that this circuit relies heavily on temporal coding, whether it’s short-term dynamics, coincidence detection or population level activity, rather than inherent target-selective circuit wiring.

## Materials & Methods

### Animals

All experiments were conducted in accordance with the National Institutes of Health (NIH) guidelines and with the approval of the New York University Grossman School of Medicine Institutional Animal Care and Use Committee. Mice were obtained from Jackson Laboratory, and subsequent breeding was established in-house. Acute slice electrophysiology experiments used CaMKII-Cre, Pvalb-Cre, SST-Cre, CCK-Cre, and VIP-Cre transgenic mice from both sexes, 8 weeks- to 5 months-old, with a C57BL/6J genetic background.

### Stereotaxic Surgery

Animals were anaesthetized with isoflurane (1.5 to 3%, inhaled) and buprenorphine (0.1 mg/kg, injected intraperitoneally). Mice were placed in a stereotaxic apparatus (Stoelting), a small incision (∼0.5 cm length) was made in the skin to expose the skull, and the skull was leveled flat according to bregma and lambda. A small craniotomy (∼0.5 mm diameter) was drilled into the skull above the injection sites on the right hemisphere and 92 to 368 nl of virus was injected (Drummond Scientific Nanoject II) into the brain at the following coordinates:

| Brain Region | LEC | LEC | MEC | MEC | CA3 | CA3 |
| --- | --- | --- | --- | --- | --- | --- |
| Axis of Injection | Co-ordinates of Injection |  |  |  |  |  |
| Medial-lateral relative to midline (x) | 4.5 | 3.2 | 3.8 | 3.9 | 2.2 | 2.0 |
| Anterior-posterior relative to bregma (y) | -4.7 | -3.0 |  |  | -1.9 | -1.7 |
| Anterior-posterior relative to sinus (y) |  |  | 0.25 | 0.3 |  |  |
| Dorsal-ventral relative to surface of the brain (z) | -2.5 | -2.5 | -1.0 | -1.0 | -1.8 | -1.8 |
| Dorsal-ventral relative to surface of the brain (z) | -2.8 | -2.8 |  |  | -2.0 | -2.0 |
| Dorsal-ventral relative to surface of the brain (z) | -3.0 | -3.0 |  |  | -2.2 | -2.2 |
| Angle of approach (degrees) | 0 | 0 | 8 | 8 | 0 | 0 |

The injection pipette was slowly lowered into the brain to the deepest targeted coordinate, left at this location for 2 minutes prior to injection. 46 to 69 nl of virus was injected in 23 nl increments spaced by 15 seconds at each z coordinate, with an additional 2-minute pause between z coordinates and a final 10 min incubation at the last (shallowest) z coordinate of each injection site before slowly retracting the injection pipette out of the brain. Mice were sutured, given Neosporin topically on the incision site, injected with 0.5-1 ml of sterile saline subcutaneously, monitored until full recovery from anesthesia was observed, and given analgesic postoperative care (buprenorphine at 0.05-0.1 mg/kg or meloxicam at 2-5 mg/kg injected subcutaneously) for 2 days. Infection sites were confirmed to be specific post hoc for all experiments by examining fluorescence from soma and axonal projections in the hippocampus and entorhinal cortex.

For *ex vivo* slice electrophysiology experiments the following adeno-associated viruses were used:

| Virus | Experiment | Source/Reference | Identifier |
| --- | --- | --- | --- |
| AAV2/5.CaMKII.Chronos.GFP | Single-labelled,<br>GABAergic<br>silencing | Neurophotonics,<br>COVF Viral Vector<br>Core<br>(RRID:SCR_016477) | 319-aav5 |
| AAV2/5.Syn.FLEX.ChrimsonR.tdTomato | Dual-labelled,<br>Crosstalk | Addgene | 62723-AAV5 |
| AAV2/5.Syn.FLEX.Chronos.GFP | Dual-labelled | Neurophotonics,<br>COVF Viral Vector<br>Core<br>(RRID:SCR_016477) | 527-aav5 |
| 2/5.Dlx56.DIO.hM4Di.mCherry | CCK-Cre<br>GABAergic<br>silencing | plasmid obtained<br>from Yingxi Lin, AAV<br>packaged by Boston<br>Children's Hospital<br>Viral Core |  |

### *Ex vivo* Electrophysiology

All slice preparation was done as previously described in Robert. V. et al and Moore. J.J. et al.

Artificial cerebrospinal fluid (ACSF) and protective dissection ACSF (dACSF) were always oxygenated with a 95% O_2_ and 5% CO_2_ mixture. Mice were deeply anaesthetized with isoflurane (5% for 5 min, inhaled) and perfused transcardially with ∼20 ml of ice-cold NMDG-based dACSF containing (in mM): NMDG 93, KCl 2.5, NaH2PO4 1.25, NaHCO3 30, HEPES 20, glucose 25, thiourea 2, Na-ascorbate 5, Na-pyruvate 3, CaCl2 0.5, MgCl2 10. Brains were then rapidly removed, cortico-hippocampal complexes were dissected out and placed upright into a custom-made agar mold in ice-cold dACSF. Then, 400 μm thick transverse slices were cut (vibratome, Leica VT1200S) at low speed (0.04 mm/s) and blade vibration amplitude (0.5 mm) in ice-cold dACSF. Slices were transferred to an immersed-type holding chamber and maintained in ACSF containing the following (in mM): NaCl 125, KCl 2.5, NaH2PO4 1.25, NaHCO3 25, glucose 22.5, Na-ascorbate 1, Na-pyruvate 3, CaCl2 2, MgCl2 1. Slices were incubated at 32°C for ∼20 min and then maintained at room temperature for at least 30 min prior to recordings.

Individual slices were transferred to a recording chamber perfused with ACSF at 3 to 5 ml/min (peristaltic pump, Watson Marlow) at 30°C (in-line heater, Warner Instruments TC-324B). Tissue was visualized under an upright microscope (Zeiss Examiner A1 or Olympus BX51WI) equipped with DIC or Dodt gradient contrast at 5x to 40x magnification with additional zoom optics 1 to 2.5x and captured by a video camera (Hamamatsu ORCA-spark or ORCA-flash4.0). The headstage connected to the recording electrodes were mounted on motorized micromanipulators (Luigs and Neumann GmbH). Patch-clamp recordings were performed with potassium- or cesium-based intracellular solution containing the following (in mM): K- or Cs-methyl sulfonate 135, KCl 5, EGTA-KOH 0.1, HEPES 10, NaCl 2, MgATP 5, Na2GTP 0.4, Na2-phosphocreatine 10, and biocytin (4 mg/ml). Gigaohm seals were formed and whole-cell recordings were obtained from CA3 pyramidal neurons soma (blind patch). Pipette resistances were 2 to 5 megaohms, series resistances were 8 to 25 megaohms. Bridge-balance was applied in current-clamp. Unless stated otherwise, the membrane potential was held at −70 mV in current-clamp. The liquid junction potential was <10 mV and not corrected for.

Dual-color optogenetic experiments were performed with Chronos- and ChrimsonR-expressed in MEC_glu_ and LEC_glu_ (interchangeably) stimulated with 1 ms 1 to 10% 470 nm and 625 nm (0.05 to 0.31 mW/mm^2^) light pulses that were empirically determined to allow spectral separation (intensity below crosstalk threshold of ChrimsonR activation by 470 nm light). Crosstalk threshold was determined using ChrimsonR-only expressed in MEC_glu_ or LEC_glu_, axons were stimulated with 1ms 1-100% 470 nm (0.05 to 3.02 mW/mm^2^) light pulses. Monosynaptic connectivity was probed in dual-color optogenetic experiments using 1ms 1-10% 470 nm and 625 nm (0.05 to 0.31 mW/mm^2^) light pulses while perfusing the slice with ACSF containing 0.2-1 µM tetrodotoxin (TTX) to prevent sodic action potential generation, and 100 µM 4-aminopyridine (4-AP) to block KV1 potassium channels. Spike probability was probed using 1ms 1-100% 470 nm and/or 625 nm light pulses.

Single-color optogenetic experiments were performed with Chronos-expressed in MEC_glu_ or LEC_glu_ stimulated with 1ms 1 to 100% 470 nm (0 to 3.02 mW/mm^2^) light pulses. All short-term plasticity experimental stimulation was performed with 20x 1 ms pulses of low-intensity 470 nm light that elicited a reliable response (<10%, 0.31 mW/mm^2^). All interneuron modulation experiments were performed with 1ms of 100% 470 nm (3.02 mW/mm^2^) light pulses and 10 µM clozapine *N*-oxide (CNO) to activate hM4D(Gi) DREADDs was bath applied via ACSF perfusion.

Data was obtained using a Multiclamp 700B amplifier (Molecular Devices), sampled at 10 khz, digitized using a Digidata 1550B AD/DA board (Molecular Devices), and acquired with the pClamp10 software (Molecular Devices). Data analysis was performed in IgorPro (Wavemetrics) and MATLAB 2024 (MathWorks) with custom-written code.

### Histology

Non *ex vivo* electrophysiology mice were deeply anaesthetized with isoflurane (5% for 5 min, inhaled) and perfused transcardially with ∼20 ml of phosphate-buffered saline (PBS) followed by ∼20 ml of 4% paraformaldehyde (PFA) in PBS. Brains were removed and fixed in 4% PFA in PBS at 4°C for at least 24 hours. Brains were washed in 0.3 M Glycine in PBS for 15 min followed by three 15-min washes in PBS. Then, 50 to 100 μm thick coronal slices were cut (vibratome, Leica VT1000S) and stored in PBS at 4°C. Similarly, samples recovered from *ex vivo* electrophysiology (400 μm thick transverse slices) were fixed in 4% PFA in PBS at 4°C for at least 24 hours, washed in 0.3 M Glycine in PBS for 15 min followed by three 15 min washes in PBS, and stored in PBS at 4°C. Tissue was permeabilized with 0.5% Triton in PBS (PBST) for twice for 20 min, blocked with 10% NGS in 0.5% PBST for 4 hours, and incubated with primary antibodies in 3% NGS 0.1% PBST overnight at 4°C. Tissue was then washed with 0.2% PBST for 15 min followed by three 30 min washes in PBS before being incubated with secondary antibodies and Streptavidin where applicable in 3% NGS 0.1% PBST for 24 hours (100 μm slices) or 48 hours (400 μm slices) at 4°C. Lastly, slices were washed five times for 15 min in PBS and mounted in Vectashield Hard Set Mounting Medium with DAPI (Vector Laboratories).

Primary antibodies were rabbit anti-RFP (1:1000, Rockland/ThermoFisher 600-401-379) and chicken anti-GFP (1:1000, Abcam no. 13970). All secondary antibodies and dyes were purchased from ThermoFisher: Alexa Fluor 555-conjugated goat anti-rabbit (1:1000, no. A21428), Alexa Fluor 488-conjugated goat anti-chicken (1:1000, no. A11039), Alexa Fluor 647-conjugated streptavidin (1:500, no. S21374), Alexa Fluor 455-conjugated Neurotrace (1:200, no. N21479).

Samples were screened by epifluorescence imaging (Olympus VS200). Relevant samples were imaged with an inverted Zeiss Axio Observer Z1 confocal microscope using 10x (air, 0.3 NA), 20x (air, 0.8 NA) or 40x (oil, 1.3 NA) objectives (Zeiss) and 405, 488, 594, and 647 nm lasers for fluorophore excitation. Images were acquired as 1024 × 1024 pixels 16 bits Z-stacks with a 5-μm (10x, 20x) or 1 μm (40x) Z-step size and tiled in X and Y as needed to cover the samples and stitched using the Zen Microscopy software (Zeiss). Further processing of confocal or epifluorescence images was done in ImageJ.

### EC projection Analysis

We used Fiji (ImageJ) to calculate the profile plots of normalized integrated intensity of the EC projections in CA3 SLM against the distance of CA3 PN (Extended Data Fig. 1), we hand-drew a straight line across the EC projections from the edge of the DG border to the CA3 SP. The analysis was performed on maximal projections of stacks of confocal images.

### *Ex Vivo* Electrophysiology Analysis

All the electrophysiology data were analyzed, and graphs were generated using IgorPro 8.0.4.2 (Wavemetrics) and MATLAB 2024 (MathWorks) using custom-written scripts. For all light-evoked postsynaptic responses, peak amplitude was measured as the response local maximum (for PSPs and IPSCs) or minimum (for EPSCs) from the baseline. The baseline value was averaged over 100 ms before stimulation onset. The summation was measured as the peak of the second response relative to the baseline of the first response. The rise time was measured between 10-90% of the time to peak. The decay was measured between 10-90% of the time to decay before the next stimulation. The latency was measured as the time from stimulation to the onset of the response. The half-width was measured as the duration of the response at 50% of its maximal amplitude. The AUC of individual responses (Extended Data Fig. 2) was measured between 10% rise-90% decay times. For current-clamp experiments (Fig. 5), any spiking was omitted from subthreshold input-output calculations. Response indices (Fig. 2) were calculated using the amplitudes of the CA3 response to individual input stimulation: MEC_glu_ response – LEC_glu_ response / MEC_glu_ response + LEC_glu_ response. All response ratios (Fig. 3, 5, Extended Data Fig. 3,4) were taken from individual recordings and were then averaged across a cell. The AUC for STP experiments (Extended Data Fig. 3) was calculated as the area under the entire response (peak to baseline) with the baseline calculated before stimulation onset. For GABAergic silencing experiments (Fig. 4), the pre- and post-silencing IPSC amplitude was determined by averaging the steady-state responses for 8 data points (2 min) before and after (∼10 min) CNO application.

### *Ex Vivo* Statistics

Results are reported ±SEM. Normality was tested with the Shapiro-Wilk and Kolmogrov-Smirnov test, and variance were compared with the Barlett test, to choose between parametric and nonparametric statistical analysis. Statistical significance was assessed using the Mann-Whitney test, paired *t* test, Wilcoxon signed-rank test, one-way ANOVA, Mixed-Effects Model, two-way ANOVA where appropriate. Geisser-Greenhouse correction was used for all Mixed-Effects Model analysis. The symbols *, **, ***, and **** denote *p* values of <0.05, <0.01, <0.001, and <0.0001 respectively. All statistics were performed in Prism (GraphPad) and figures were compiled using MATLAB 2024 (MathWorks) and Adobe Illustrator 2026.

### *In vivo* Electrophysiology Analysis

To assess CA3 population response to EC inputs (Fig.6, Extended Data Fig. 5), we used a published dataset in which current source density source-localization identified dentate spikes (DS) originating in either LEC (DS_LEC_) or MEC (DS_MEC_). From this dataset, we selected all recordings in which at least 1 single unit was localized in hippocampal area CA3, yielding 257 units from 22x 15-minute recording sessions in 3 mice (n=16, 53, 188 units per mouse; n=2-41 units per session, mean 11.68, std 8.64; 3, 6, 13 sessions per mouse). For each type of dentate spike (or combination of dentate spikes, for combination analysis), we calculated a 1-second peri-stimulus time histogram (PSTH) centered on the DS event (or second DS in the combination). The PSTH for each unit was defined as the probability of observing at least one spike from that unit within each 10-ms window within the 1-second interval, using a sliding window with step size 2 ms. We quantified the response of each unit by z-scoring the probability values of its PSTH and assessing the max, or difference between the max and min, z-score observed over a 40-ms wide window centered on the time point of maximal population response. The time point of maximal population response was identified as t= −4 ms, for both DS_MEC_ and DS_LEC_ events, identified by inspection of the population-averaged PSTHs. PSTH probabilities were z-scored to normalize for individual differences in the level of background variability in excitability per unit, but we ran the same statistical comparisons without using z-scoring (i.e. on the raw probability trace itself) and reached the same conclusions.

## Author Contributions

K.O., V.R., J.B. conceived the project and designed experiments; J.B. secured funding and supervised the project and personnel; K.O. performed all stereotaxic surgeries, histology, *ex vivo* electrophysiology experiments, data analysis, statistical tests, and coordinated the project; V.R. assisted with stereotaxic injections, and acute slice electrophysiology related to dual-optogenetic implementation and LEC_glu_ to CA3 GABAergic silencing experiments, V.R. assisted with *ex vivo* analysis throughout; L.A.A. and A.A.F. designed *in vivo* data analysis using experimental data collected under the supervision of A.A.F.; L.A.A. performed all *in vivo* analysis; B.K. performed confocal imaging and collected anatomical fluorescence data; K.O. generated all graphs, prepared all figures for visualization with input from V.R., L.A., A.F., and J.B.; L.A. and K.O. prepared *in vivo* figures; K.O., V.R., L.A.A., A.A.F., and J.B. wrote the manuscript.

## Acknowledgements

This work was supported by the following grants to J.B.: National Institutes of Health (NIH) National Institute of Neurological Disorders and Stroke (NINDS) BRAIN Initiative R01NS109994, NIH NINDS R01NS109362, NIH NINDS RM1NS132981, NIH National Institute of Mental Health (NIMH) R01MH122391, NIH National Institute on Aging (NIA) R01AG094086; Alzheimer’s Association AARGD-NTF-23-1151101, McKnight Scholar Award in Neuroscience, Klingenstein-Simons Fellowship Award in Neuroscience, Alfred P. Sloan Research Fellowship, Mathers Charitable Foundation Award, Whitehall Research Grant, a Blas Frangione Young Investigator Research Grant, New York University Whitehead Fellowship for Junior Faculty in Biomedical and Biological Sciences, and Leon Levy Foundation Award. K.O. was supported by an NIH 5T32MH019524-30 training award; V.R. was supported by a Young Researchers Bettencourt Prize, B.K. was supported by a McKnight Endowment Fund for Neuroscience Mathew Pecot URM Award and an NIH NINDS 5R25NS125593 BRAIN Training grant (M. Minen, NYU GsoM). A.A.F. and L.A.A. were supported by NIH NIMH R01MH132204 and NIH NIMH R01MH137669.

We are grateful for Y. Lin and M. Meng for coordinating and gifting virus for testing and the plasmid used to make virus for the CCK GABAergic silencing experiments and was commercially generated by the Boston Children’s Hospital Viral Core, which is supported by NIH5P30EY012196. We thank P. Frazel for plasmid preparation. We are grateful to O. Bilash, T. Butola, M. Druart, M. Hopkins, J. Moore, K. Nagel, N. Nambiar, S. Peron, and R. Tsien for their input at various stages of the project.

## Competing Interests

### Materials & Correspondence

Materials requests to Jayeeta Basu Correspondence to Vincent Robert & Jayeeta Basu

### Data Availability

Data and custom code available upon request to Jayeeta Basu and André Fenton.

---

**Extended Data Figure 1:**
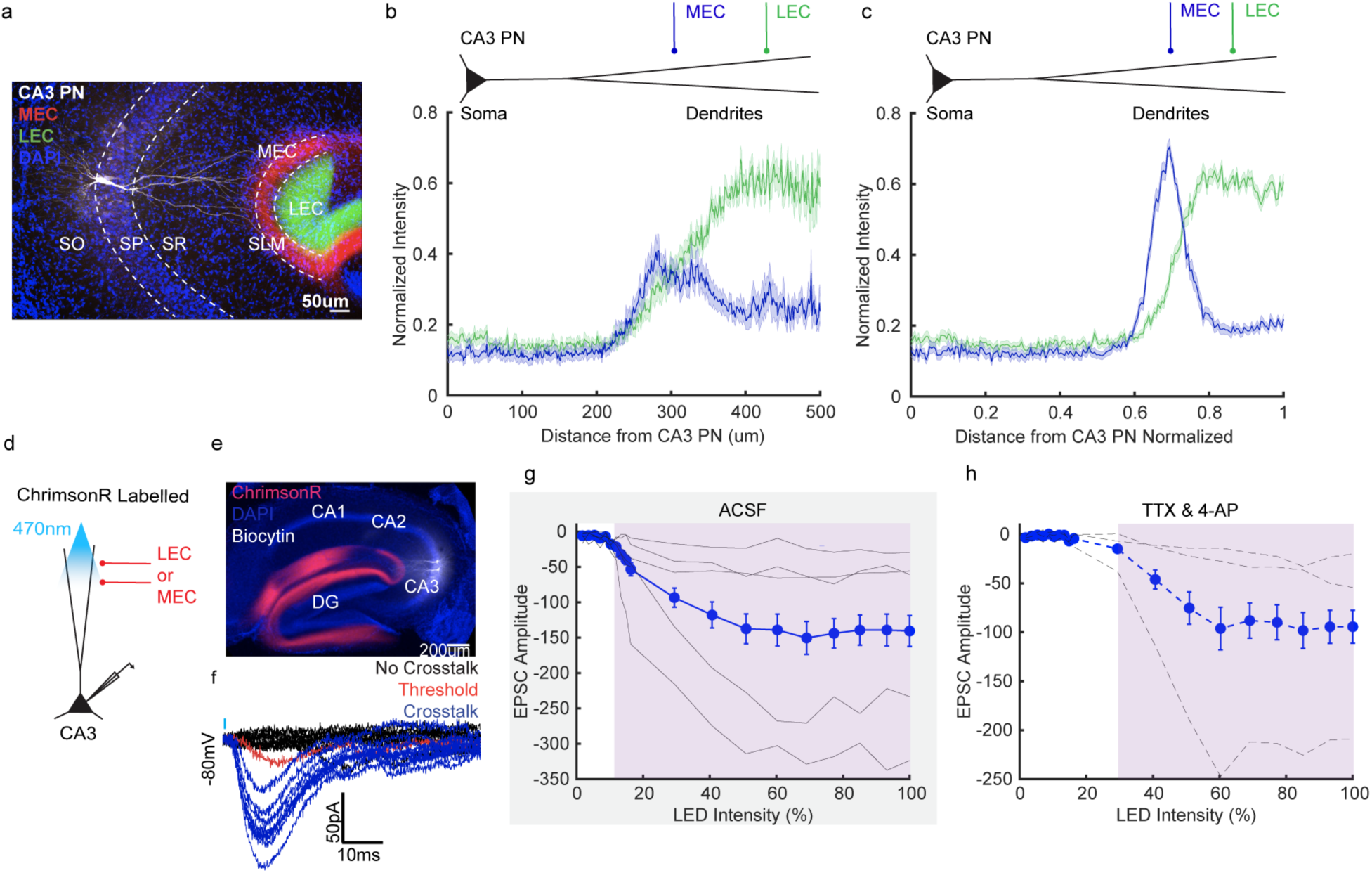
**a,** Confocal image of CA3 region of 400 µm transverse hippocampal, mouse brain slice showing recorded CA3 a/b PN filled with Biocytin (white) and MEC axons (red) and LEC axons (green). White dashed lines denote where measurements for intensity were taken from. **b,c** Normalized fluorescence intensity of MEC (blue) and LEC (green) axons CA3 region of the hippocampus as a function of distance from the CA3 PN layer (SP) with **b,** and without **c,** normalization to the PN layer (n = 32 slices, N = 16 mice). The bold line is mean fluorescence intensity and the shaded area represents SEM. **d,** Schematic of ChrimsonR single-labelled viral injection strategy. AAV2/5.Syn.flex.ChrimsonR.tdTomato was injected into MEC or LEC of CaMKII-Cre mice to drive excitatory opsin expression selectively in glutamatergic neurons. To test for crosstalk, we recorded responses of CA3PN voltage-clamped at −80 mV to stimulation using 470 nm (blue) light over the axons. **e,** Confocal image of transverse hippocampal recording slices with MEC_glu_-(red) axons labelled with ChrimsonR. **f,** Sample traces of blue light (470 nm) ChrimsonR-driven CA3 PN EPSCs (−80 mV) at a range, 1-100% (0.05-3.02 mW/mm^2^), of LED intensities. Traces that have no time-locked response to blue light stimulation are in black (No Crosstalk), traces that have a variable time-locked response to blue light stimulation are labelled in red (Threshold), and traces that have time-locked response to blue light stimulation are in blue (Crosstalk). **g,h** Input-output function curves of light-evoked CA3 PN EPSC amplitudes across all light intensities in **g,** ACSF and **h,** 1 µM TTX & 10 µM 4-AP (1-100%). The purple area demarcates when the average response of the ChrimsonR-labelled axons to blue light is consistent, aka crosstalk (f: n = 6 cells, N = 3 mice g: n = 3 cells, N = 2 mice).

**Extended Data Figure 2:**
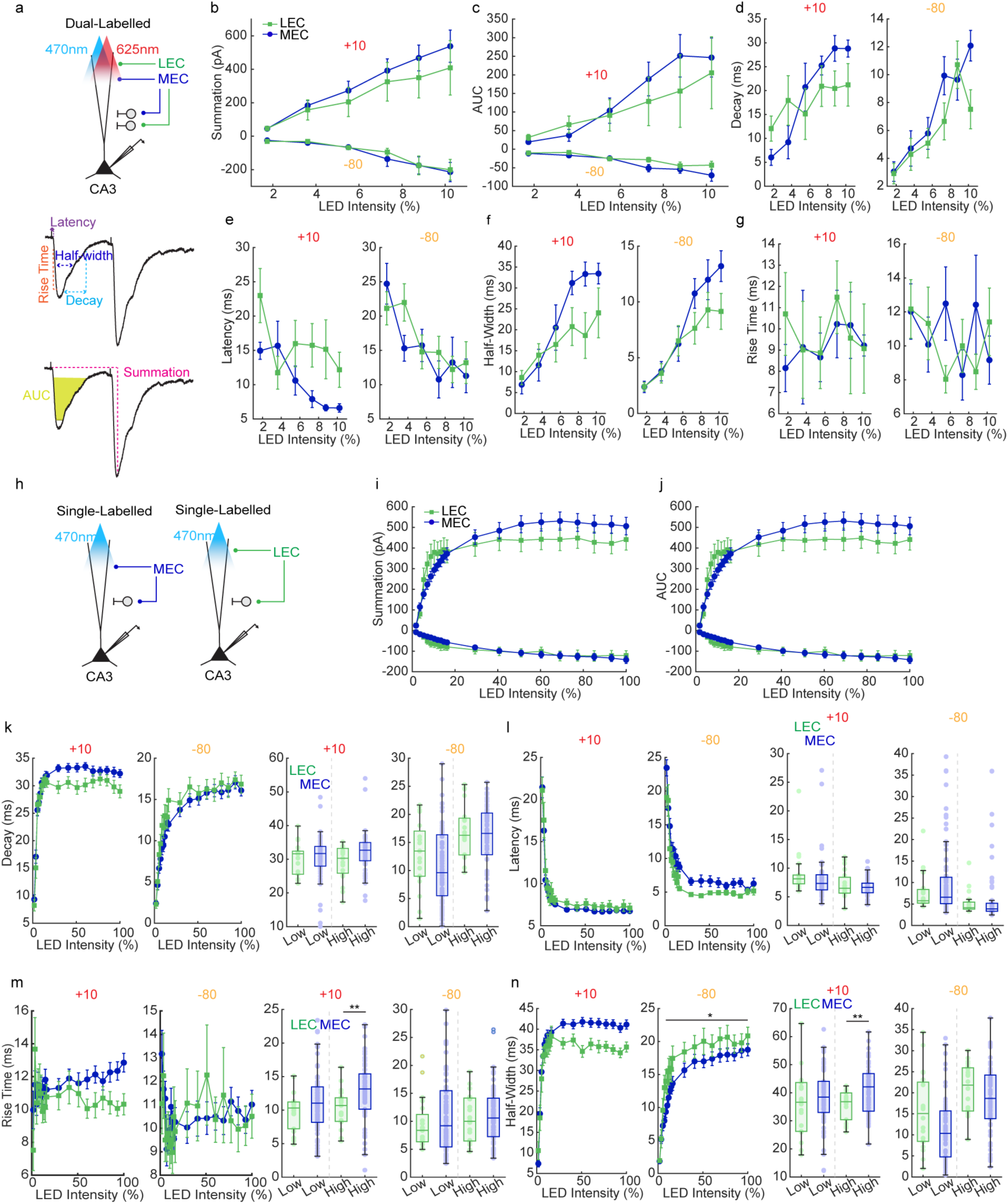
**a,** Top: Dual-labelled experimental strategy used in Figure 1a-d and Figure 2a-h. We recorded responses of CA3 PN voltage-clamped at +10 mV and −80 mV to LEC_glu_ (green) and MEC_glu_ (blue) stimulation done using 470 nm and 625 nm light over the SLM region of CA3. Stimulation was 2x 1 ms LED pulses 100 ms apart. Middle/Bottom: Schematic of PSC kinetic properties measured. **b-g**, Input-output function curves comparing the IPSC (+10 mV) and EPSC (−80 mV) properties for MEC_glu_ (blue) and LEC_glu_ (green): **b**, summation of second response **c**, area-under-the-curve (AUC) **d**, 20-80% decay **e,** latency to PSC onset **f,** duration at half-maximum (half-width) **g,** 20-80% rise time (IPSC: n = 6 cells, N = 4 mice; EPSC: n = 16 cells, N = 8 mice). All statistics are listed in Table 1. **h,** Single-labelled experimental strategy used in Figure 2i-n. We recorded responses of CA3 PN voltage-clamped at +10 mV and −80 mV to either MEC_glu_ (blue) or LEC_glu_ (green) stimulation done using 470 nm light over the SLM region of CA3. Stimulation was the same as described in **a**. **i-n**, Input-output function curves (left) and summary plots (right) comparing the IPSC (+10 mV) and EPSC (−80 mV) properties for MEC_glu_ (blue) and LEC_glu_ (green): **i,** summation of second response **j,** area-under-the-curve (AUC) **k,** 20-80% decay **l,** latency to PSC onset **m**, duration at half-maximum (half-width) **n,** and 20-80% rise time. All error bars in the input-output curves represent SEM. In the summary box-and-whisker plots, the bold line inside the box is the median, the box represents the lower and upper quartiles, and the whiskers represent the minimum and maximum values that are not outliers. All individual data points are the averaged responses of a cell to a single low or high light intensity (IPSC MEC_glu_: n = 63 cells, N = 30 mice, LEC_glu_: n = 18 cells, N = 11 mice, EPSC MEC_glu_: n = 72 cells, N = 32 mice, LEC_glu_: n = 20 cells, N = 12 mice). All input/output curve statistics are in Table 1 and high/low comparisons are in Table 2.

**Extended Data Figure 3:**
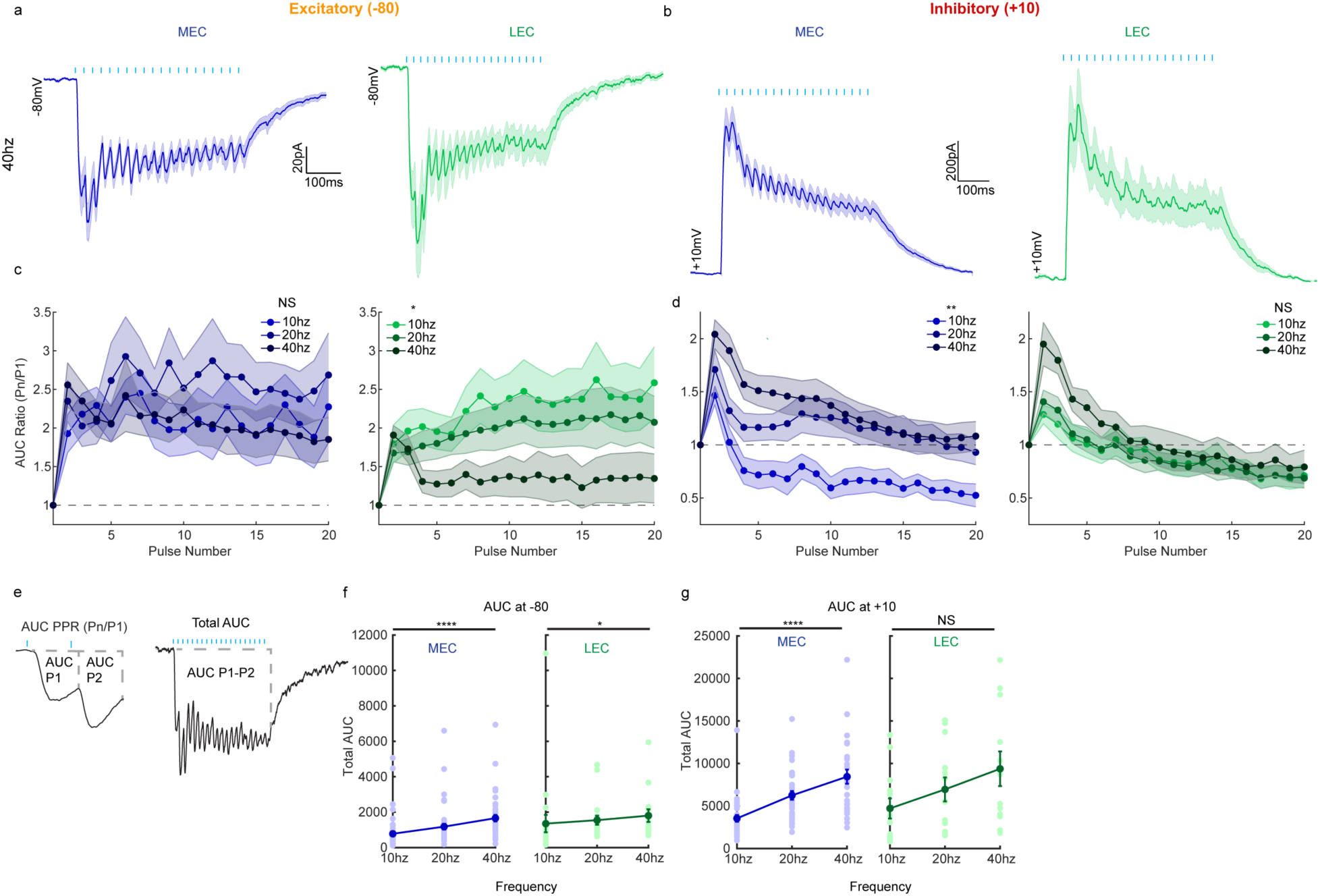
**a,b** Averaged traces of CA3 PN responses to 40 hz LEC_glu_ (green) and MEC_glu_ (blue) stimulation in voltage-clamp at −80 mV (**a**) and at +10 mV (**b**) **c, d,** Summary plots of ratio of the AUC for each stimulation to the AUC for stimulation 1 (P_n_/P_1_) of the 1-20th MEC_glu_ (blue, right) and LEC_glu_ (green, left) driven EPSCs (**c**) and IPSCs (**d**) at 10, 20, and 40 hz (MEC_glu_: EPSC NS p=0.1337, IPSC **p= 0.0021 Mixed-Effects Model w/ Geisser-Greenhouse correction; LEC_glu_: EPSC *p=0.0127, IPSC NS p = 0.1093 Mixed-Effects w/ Geisser-Greenhouse correction). **e,** Schematic of AUC properties measured, AUC PPR (left) and AUC from stimulation 1-20 (right) **f,g** Total AUC for MEC_glu_ (blue, left) and LEC_glu_ (green, right) at 10, 20, and 40 hz at −80 mV (**f,** MEC_glu_ ****p<0.0001 one-way ANOVA; LEC_glu_ *p = 0.0277 one-way ANOVA) and +10 mV (**g,** MEC_glu_ ****p<0.0001 one-way ANOVA; LEC_glu_ NS p=0.0791 one-way ANOVA). Each data point represents the mean total AUC (sum of AUC of P1-P20) of an individual cell. All error bars represent SEM. (MEC_glu_ −80 mV 10 hz: n = 54 cells, N = 27 mice, 20 hz: n = 54 cells, N = 28 mice, 40 hz: n = 51 cells, N = 27 mice; LEC_glu_ −80 mV 10 hz: n = 23 cells, N = 13 mice, 20 hz: n = 20 cells, N = 12 mice, 40 hz: n = 16 cells, N = 9 mice; MEC_glu_ +10 mV 10 hz: n = 35 cells, N = 19 mice, 20 hz: n = 33 cells, N = 18 mice, 40 hz: n = 30 cells, N = 18 mice; LEC_glu_ +10 mV 10 hz: n = 14 cells, N = 9 mice, 20 hz: n = 14 cells, N = 9 mice, 40 hz: 13 cells, N = 9 mice)

**Extended Data Figure 4:**
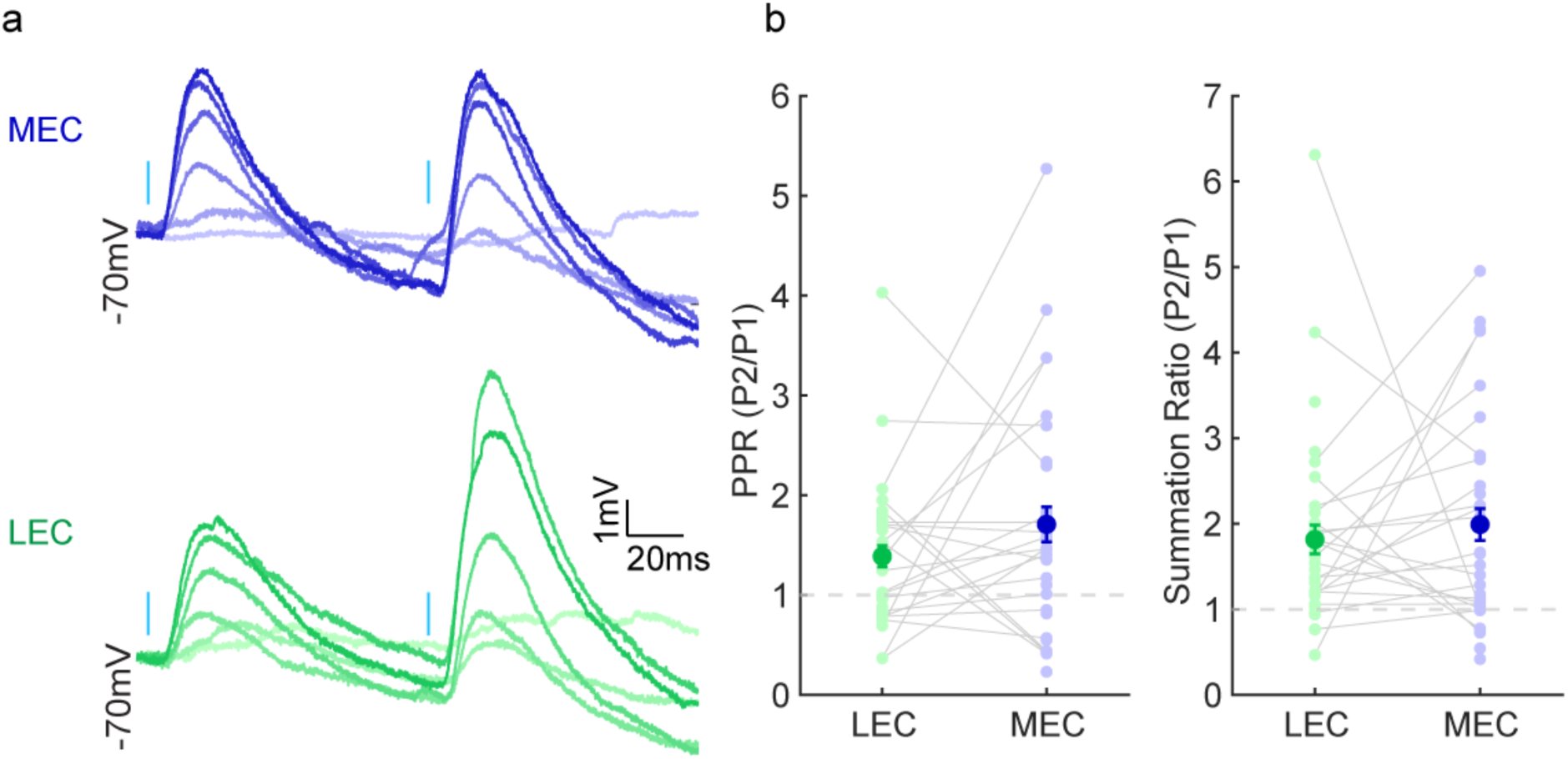
**a,** Sample traces of MEC_glu_ (blue) or LEC_glu_ (green) light-evoked CA3 PN PSPs at 10 hz stimulation at −70 mV across a range of low (0-10%) light intensities. Responses to MEC_glu_ and LEC_glu_ are from the same cell/animal. Saturation of trace indicates LED intensity. **b,** Left, summary data of paired-pulse ratio (PPR) of MEC_glu_- (blue) and LEC_glu_- driven (green) CA3 PN PSPs at 10 hz. Each data point represents average PPR from an individual cell (MEC_glu_: n = 29 cells, N = 14 mice; LEC_glu_: n = 31 cells, N = 16 mice). Right, summary data of summation ratio of MEC_glu_- and LEC_glu_-driven PSPs at 10 hz. Each data point represents the average summation ratio from an individual cell (MEC_glu_: n = 29 cells, N = 14 mice; LEC_glu_: n = 31 cells, N = 16 mice).

**Extended Data Figure 5:**
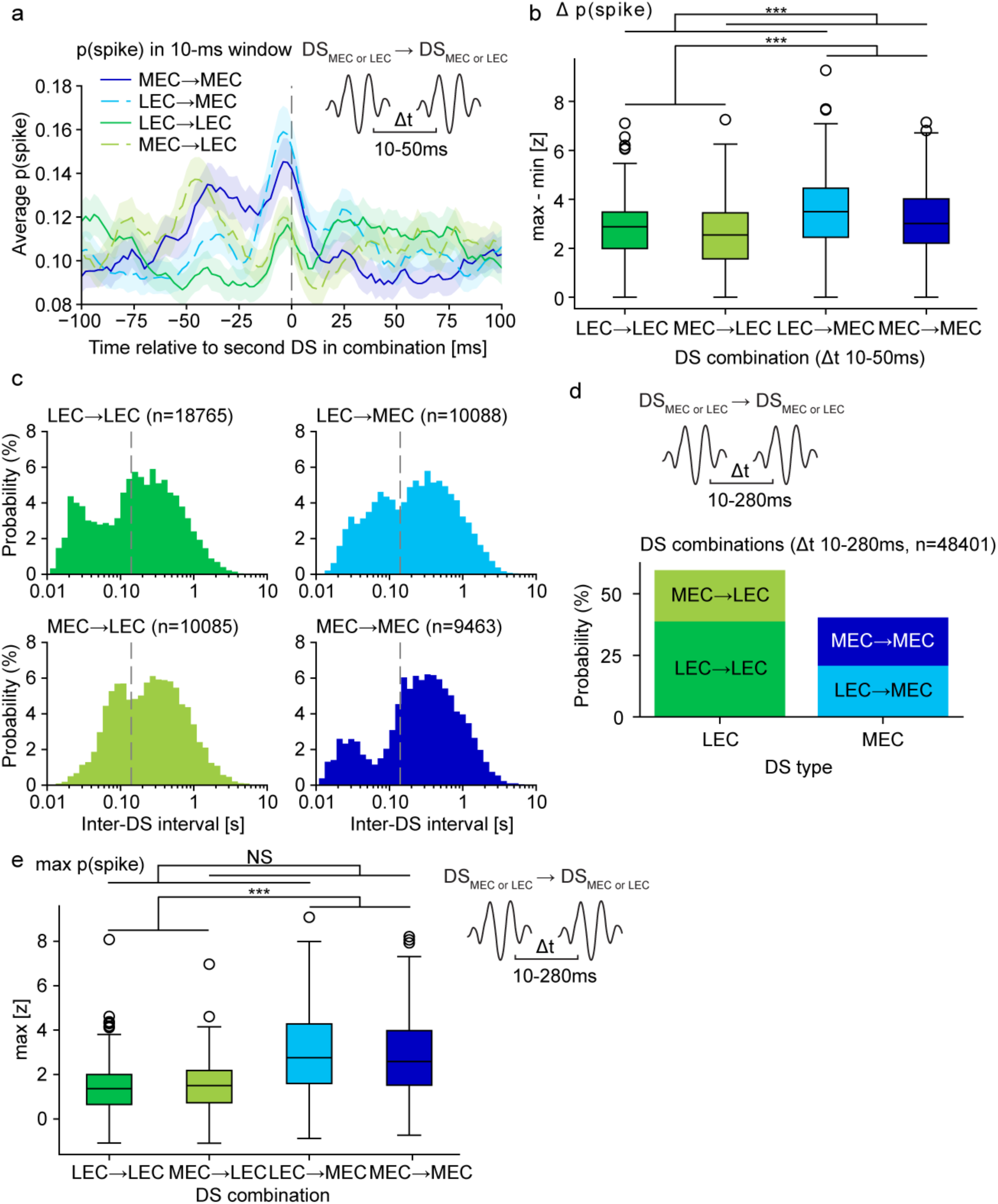
**a,** Same as Fig. 6e, for DS event combinations with inter-DS intervals in the range 10-50 ms. **b,** Same as Fig. 6f, for DS event combinations with inter-DS intervals in the range 10-50 ms. A two-way ANOVA was used to evaluate differences in response strength as described in Fig. 6f. Current DS: F1,256 = 61.205, ***p = 0.00; preceding DS: F1,256 = 22.728, ***p = 0.00; interaction: F1,256 = 0.0007, NS p = 0.980. Using max p(spike) scoring (as described in Fig. 6c, left), current DS: F1,256 = 91.576, ***p = 0.00; preceding DS: F1,256 = 3.491, NS p = 0.0628; interaction: F1,256 = 2.549, NS p = 0.112. **c,** Distributions of interspike intervals for the four DS event combinations. **d,** Probability of each of the four DS event combinations. Binomial test of proportions within each of LEC and MEC given the overall LEC/MEC ratio, LEC: z = 18.81, ***p = 0.00, MEC: z= 22.83, ***p = 0.00. **e,** Same as Fig. 6f, if response is quantified using max z-scored p(spike), as described in Fig. 6c (left). A two-way ANOVA was used to evaluate differences in response strength as described in Fig. 6f. Current DS: F1,256 = 124.61, ***p = 0.00; preceding DS: F1,256 = 2.328, NS p = 0.128; interaction: F1,256 = 3.254, NS p = 0.0724.

